# Lipid droplets restrict phagosome formation during *Candida* challenge

**DOI:** 10.1101/2024.06.11.598578

**Authors:** Wanwei Sun, Han Wu, Guimin Zhao, Qing Shui, Lei Zhang, Xiaoxi Luan, Tian Chen, Feng Liu, Yi Zheng, Wei Zhao, Xiaopeng Qi, Bingyu Liu, Chengjiang Gao

## Abstract

Lipid droplets (LDs) are intracellular organelles which can be induced and interact with phagosomes during the process of pathogen phagocytosis in macrophages. However, the function of LDs in the phagocytosis remains elusive. Here, we unveil the role of LDs in modulating phagosome formation using a fungal infection model. Specifically, LDs accumulation restricted the quantity of phagosome formation, and protected macrophages from death. Mechanistically, LDs formation competitively consumed intracellular endoplasmic reticulum membrane and altered RAC1 translocation and its GTPase activity, which resulted in limiting phagosomes formation in macrophages during fungi engulfed. Mice with *Hilpda*-deficient macrophages were more susceptible to the lethal sequelae of systemic infection with *C. albicans*. Notably, administration of ATGL inhibitor Atglistatin improved host outcomes in disseminated fungal infections. Taken together, our study elucidates the mechanism of LDs in controlling phagosomes formation to avoid immune cell death, and offers a potential drug target for treatment of *C. albicans* infections.

**Highlights:**

- *C. albicans* infection induces triglycerides production and LD accumulation in macrophages.
- LD restricts the quantity of phagosome formation and protects macrophages from death.
- LD consumes intracellular ER membrane components and alters RAC1 translocation and its GTPase activity.
- Administration of ATGL inhibitor Atglistatin improves host outcomes in disseminated fungal infections.

## INTRODUCTION

*Candida albicans* are the most frequently isolated human fungal pathogens and have become one of the leading causes of infection in hospital settings.^1^ Each year, invasive fungal infections lead to ~700,000 cases with approximately 40% mortality.^2,3^ Currently, the available antifungal drugs for systemic candidiasis are limited, such as azoles, echinocandins and polyenes.^4,5^ The widespread drug resistance and drug toxicity further restrict the application and lead to high mortality rate.^6,7^ As such, a deeper understanding of host antifungal immunity is crucial for the development of novel drugs and improving our future therapeutic strategies.

Upon fungal infection, the pro-inflammatory chemokines and cytokines induced by the C-type lectin receptors associated signaling trigger neutrophil infiltration, macrophage maturation, and T cell differentiation including T helper-1 (Th1) and T helper-17 (Th17) subsets.^8,9^ The phagocyte-dependent phagocytosis is critical for clearance and killing fungal pathogens during systemic candidiasis.^10,11^ Phagocytosis is initiated by the binding of the polysaccharide component of fungal cell wall to Dectin receptors at the cell surface.^12^ Signaling events rapidly lead to cytoskeletal (mainly actin) reorganization and concomitant pseudopodia extension.^13^ The enrichment of PI (4,5) P_2_ is critical for activation of small GTPases of the Rho family (RAC and CDC42) to induce actin polymerization, which generates the force driving membrane extension.^14^ Actin continues to be actively polymerized to extend the phagosomal membrane, while being remodeled at the base of the phagocytic cup, allowing intracellular membrane recruitment.^14^ Further actin polymerization and contractile activity lead to membrane apposition and eventually fusion, which is catalyzed by the scission activity of Dynamin 2.^14^ Although lysosomes and recycling endosomes have been shown to be able to supply membrane during phagocytosis, the ER (endoplasmic reticulum) is likely to be the main source of intracellular membrane contributing to phagosome formation in macrophages.^12^ After their formation, phagosomes fuse sequentially with early endosomes, late endosomes and lysosomes.^15,16^ Our recent study also finds that the endoplasmic reticulum protein STING translocates along with ER membrane to phagosomes to negatively regulate anti-fungal immunity.^17^ After engulfed by phagocytes, *C. albicans* can escape from immune cells through forming hyphae and make cells death. The greater the number of *C. albicans* engulfed by phagocytes, the quicker the phagocytes perish.^18,19^ However, the mechanisms by which phagocytes use to control the number of phagosomes to avoid phagocytes death caused by excessive engulfment, has not be reported so far.

LD (Lipid droplets) is a lipid rich intracellular organelle derived from ER and wrapped by phospholipid monolayer. Their surfaces are attached with a variety of structural proteins. The core neutral lipid components are mainly triglycerides and cholesteryl ester.^20–22^ LD plays a crucial role in cell metabolism, lipid biology, and cell signal transduction.^23^ The accumulation of intracellular LD depends on the balance between lipid synthesis and degradation, and plays a key regulatory role in energy supply and redox homeostasis.^20^ The *De novo* synthesis of triglycerides is mainly carried out by GPAT (Glycerol-3-phosphate O-acyltransferase), AGPAT (1-acylglycerol-3-phosphate O-acyltransferase), PAP (Phosphatidic acid phosphatase) and DGAT (Diacylglycerol acyltransferase). In four consecutive reactions catalyzed by them, one molecule of 3-phosphoglycerol and three molecules of fatty acid are synthesized into one molecule of triglycerides. Meanwhile, fatty acids in triglycerides are hydrolyzed sequentially by ATGL (Adipose triglyceride lipase), HSL (Hormone sensitive lipase) and MAGL (monoacylglycerol lipase). The fatty acids generated by hydrolysis can be oxidized in mitochondria, providing metabolic energy for cell survival in malnutrition environment.^22^ HILPDA (hypoxia-inducible lipid droplet-associated) is an inhibitor of macrophage lipolysis, which can promote the accumulation of LD by inhibiting the activity of ATGL.^21^ The protein compositions associated with LD varies, depending on the metabolic state and stimulation conditions.^24^ In addition, LD is involved in the pathogenesis of various pathogens, including viruses, parasites and bacteria, and plays multiple roles in mediating cellular inflammation, immune metabolism, and host pathogen interactions.^25^ A recent study shows that LD can drive or assist the innate immune system in defending against *Escherichia coli* (*E. coli*), and the LD assembles and organizes multiple host defense proteins, including interferon-inducible guanosine triphosphatases and the antimicrobial cathelicidin, to kill intracellular pathogens.^26^ However, whether LD can protect the host in the process of *C. albicans* infection need to be explored.

Here, we studied the mechanisms as to how macrophages control excessive phagosomes formation to protect its survival. We discovered that LD functions as a negative regulator to modulate the phagocytosis through competitive consumption of intracellular ER membrane components and alteration of RAC1 GTPase activity upon *C. albicans* stimulation. Notably, we found that administration of inhibitors of ATGL, namely Atglistatin, protected mice from the lethal sequelae of systemic *C. albicans* infections. Collectively, our results defined how the interplay between host lipid metabolism and phagocytosis of fungal pathogens determines the outcomes of immune protections and disease in systemic candidiasis.

## RESULTS

### Fungal stimulation induces triglycerides production and LD accumulation

To determine whether the metabolites were involved in anti-fungal immunity, we performed a metabolomics profiling to analyze the changed metabolites in BMDMs (bone marrow-derived macrophages) stimulated with fungal component D-zymosan (Zymosan Depleted, Dectin-1 ligand). We identified 72 differential metabolites, among which 42 were up-regulated and 30 were down-regulated in D-zymosan stimulation group, compared to that in control group (Figure S1A). Principal component analysis showed a distinct separation of global metabolic expression profile based on D-zymosan stimulation (Figure S1B). Among the 42 up-regulated metabolites, there were 27 triglycerides with different branched chains (Figure 1A and S1C). KEGG enrichment analysis revealed that these metabolites were involved in thermogenesis, regulation of lipolysis in adipocytes, fat digestion and absorption, glycerolipid and cholesterol metabolism (Figure S1D).

**Figure 1.**
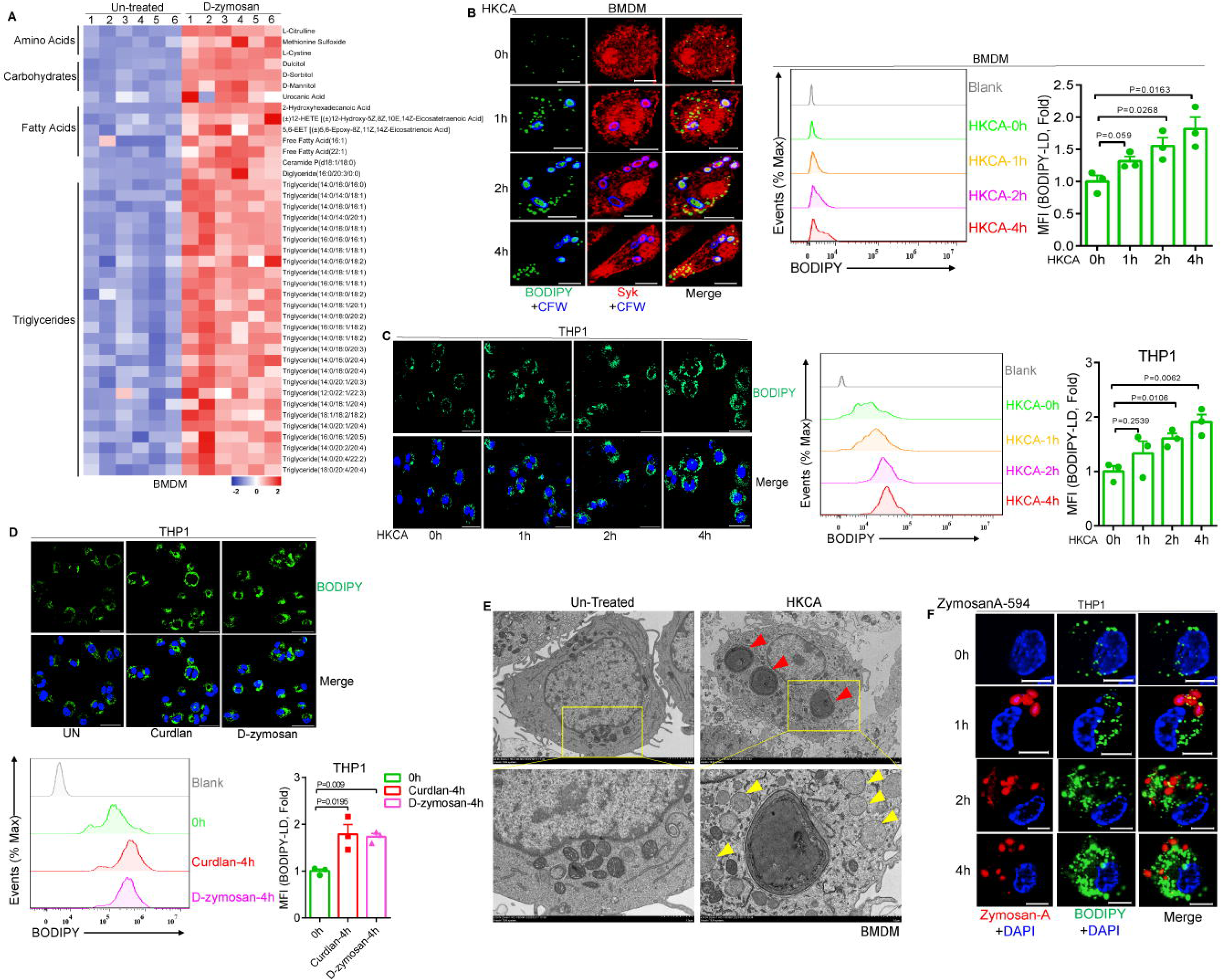
Fungal stimulation induced triglycerides production and LD accumulation. **(A)** Heatmap of changes in metabolites of selected up-regulated metabolites in D-zymosan-stimulated mice bone marrow derived macrophages (BMDMs) versus non-stimulated macrophages. All differential metabolites are displayed in FigureS1A and Table S1. All identified metabolites are shown in Table S2. **(B)** BMDMs were stimulated with HKCA (MOI=3) for the indicated times followed by immunofluorescence staining. LD, HKCA and cell was stained by BODIPY, CFW and Syk, respectively (Scale bar: 10μm). Meanwhile, LD-BODIPY MFI was measured at the indicated times by flow cytometry. Error bars represent SEM of three biological replicates. **(C)** THP1 cells were stimulated with HKCA (MOI=3) for the indicated times followed by immunofluorescence staining. LD was stained by BODIPY and nucleus was stained by DAPI. (Scale bar: 40μm). Meanwhile, LD-BODIPY MFI was measured at the indicated times by flow cytometry. Error bars represent SEM of three biological replicates. **(D)** THP1 cells were stimulated with Curdlan (100μg/mL) or D-zymosan (100μg/mL) for 4h followed by immunofluorescence staining. LD was stained by BODIPY and nucleus was stained by DAPI. (Scale bar: 40μm). Meanwhile, LD-BODIPY MFI was measured by flow cytometry. Error bars represent SEM of three biological replicates. **(E)** Electron microscopy images of a non-stimulated and stimulated with HKCA (MOI=3) for 4h BMDM. LDs in the HKCA-stimulated BMDM are indicated by yellow arrowheads. Phagosomes are indicated by red arrowheads. **(F)** THP1 cells were stimulated with ZymosanA-Alexa Fluor™ 594 (100μg/mL) for the indicated times followed by immunofluorescence staining. LD was stained by BODIPY and nucleus was stained by DAPI. (Scale bar: 10μm). Two-tailed unpaired Student’s t test (D) and one-way ANOVA for (B and C).

We then utilized TLC (thin layer chromatography) assay to validate the change of intracellular triglycerides level. Consistent with the metabolomics, triglycerides level was obviously increased after BMDMs were stimulated with D-zymosan (Figure S1E). Similarly, triglycerides level was also increased in BMDMs stimulated with HKCA (Heat-killed *Candida albicans*) or Curdlan (another Dectin-1 ligand) (Figure S1E). We further quantified the triglycerides level using triglyceride test kit and the results showed triglycerides level was significantly up-regulated in BMDMs stimulated with D-zymosan, Curdlan or HKCA (Figure S1E). The triglycerides level was also increased in RAW264.7 macrophages stimulated with D-zymosan (Figure S1F) and human monocyte THP1 cells stimulated with D-zymosan or HKCA (Figure S1G).

Triglycerides synthesized within cells are mainly stored in lipid droplets (LDs).^18^ Thus, we next examined LD formation under confocal microscopy using boron-dipyrromethene (BODIPY) to label LD. As the stimulation time extended, HKCA-induced triglycerides production was gradually increased in BMDMs (Figure 1B). Meanwhile, we quantified LDs in BMDM cells labeled with BODIPY by flow cytometry. HKCA-elicited BMDM cells displayed markedly increased staining with BODIPY, indicative of increased LD formation, when compared to untreated cells (Figure 1B). In addition, triglyceride test kit found triglycerides level was gradually expanded in BMDM cells upon HKCA stimulation (Figure S1H). Consistently, these experiments were conducted in THP1 cells and the results showed that HKCA could also induce LD formation in THP1 (Figure 1C and S1I). Besides, BODIPY-labeled LDs were also increased in THP1 cells stimulated with Curdlan or D-zymosan (Figure 1D). Furthermore, the images of transmission electron microscopy showed that the LDs were produced in BMDMs stimulated with HKCA for 4h (Figure 1E). To exclude the possibility of BODIPY staining on the ligands or HKCA itself, we utilized Alexa Fluor^TM^ 594-labeled ZymosanA to stimulate THP1 cells for different times and confocal microscopy showed that intracellular BODIPY-labeled LDs were similarly increased upon ZymosanA stimulation. Notably BODIPY-labeled LDs were not colocalized with ZymosanA (Figure 1F). Collectively, these data suggested that fungal infection could induce triglycerides production and LD accumulation in macrophages.

### Fungal stimulation induces LD accumulation via promoting lipid biosynthesis and inhibiting lipolysis

The accumulation of intracellular LD depends on the balance between lipid synthesis and degradation.^20^ To investigate the mechanisms underlying LD accumulation in macrophages upon fungal stimulation, we performed RNA-seq (RNA sequencing) of BMDMs stimulated with curdlan for different times. Differentially expressed genes were provided in Table S3 and Table S4. A total of 807 and 1463 genes were differentially expressed at the 3h and 6 h time points, respectively, in BMDMs challenged with curdlan, relative to BMDMs without stimulation. Previous studies have reported that *C. albicans* challenge could up-regulate glycolysis and glucose import, but oxidative phosphorylation, mitochondrial biogenesis and activity were turned down.^18^ Consistent with the report, we also observed that curdlan stimulation triggered upregulation of the genes involved in glycolysis and the concomitant repression of genes involved in oxidative phosphorylation in mitochondrial (Figure 2A). Next, we analyzed the transcript level of lipid metabolism related genes. We found curdlan stimulation promoted the expression of genes related to lipid biosynthesis such as diacylglycerol acyltransferase-1 (*Dgat1*), and inhibited the expression of genes involved in lipolysis such as adipose triglyceride lipase (ATGL), hormone-sensitive lipase (HSL), monoacylglycerol lipase (MAGL) (Figure 2A). Notably, curdlan stimulation increased the expression of hypoxia-inducible lipid droplet-associated (HILPDA), which is an inhibitor of ATGL. However, the expression of genes involved in lipid uptake were not consistent (Figure 2A and S2B). We also analyzed the RNA-seq data reported by Timothy et al. (2018) in BMDMs infected with live *Candida albicans* and found that the genes of lipid biosynthesis were similarly increased (especially the gene of HILPDA) and the genes of lipolysis was suppressed (Figure S2A).^18^

**Figure 2.**
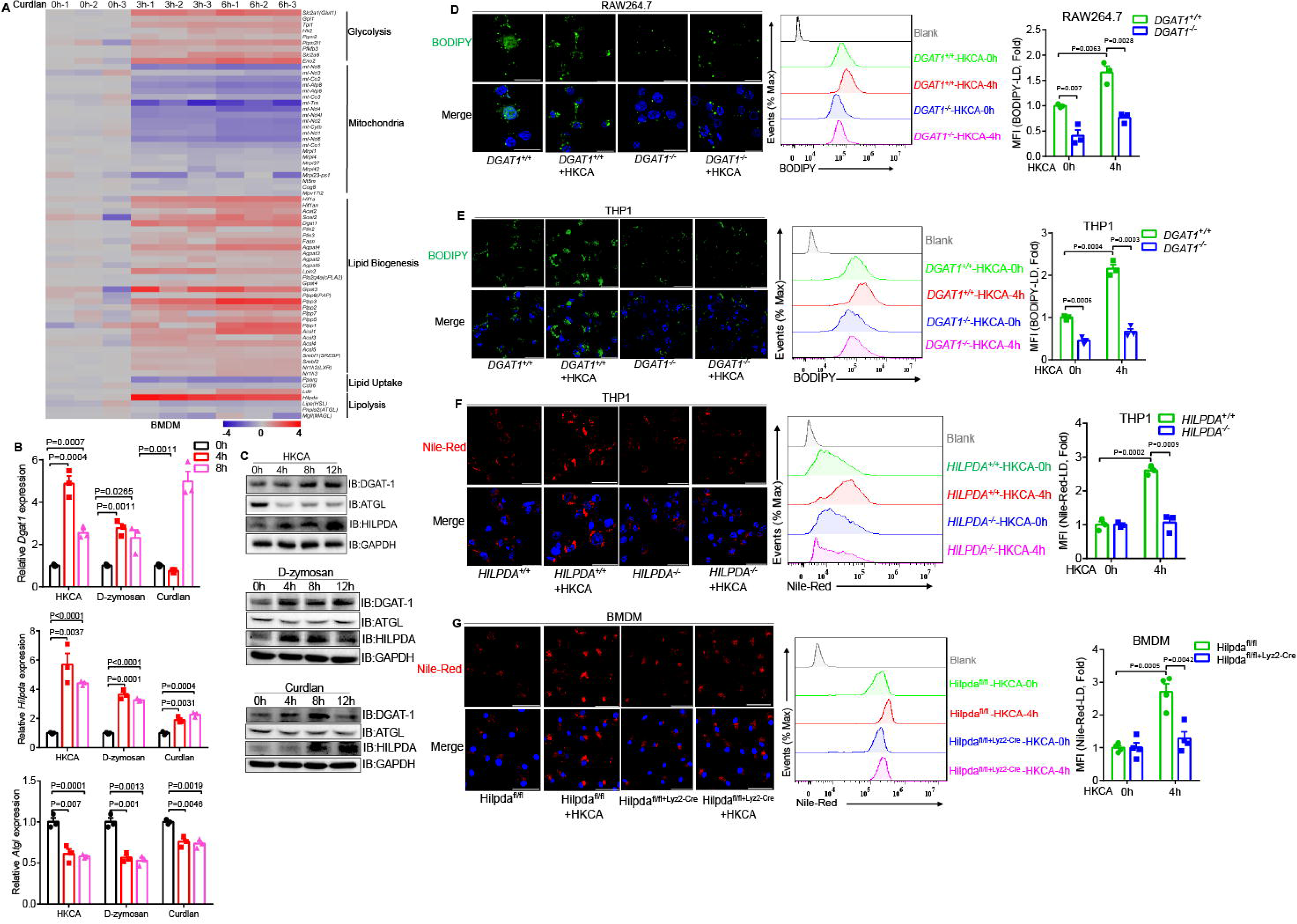
Fungal induced LD accumulation through promoting lipid biosynthesis and inhibiting lipolysis. **(A)** Heatmap of changes in gene expression of selected genes in curdlan-stimulated BMDMs over non-stimulated BMDMs. **(B)** BMDMs were stimulated with HKCA (MOI=3), D-zymosan (100μg/mL) and curdlan (100μg/mL) for the indicated times followed by real-time PCR analysis of *Dgat1*, *Atgl* and *Hilpda* genes. Error bars represent SEM of three biological replicates. Other genes’ expression was displayed in FigureS2B. The entire list of differentially expressed genes are shown in TableS3 and TableS4. **(C)** BMDMs were stimulated with HKCA (MOI=3), D-zymosan (100μg/mL) and curdlan (100μg/mL) for the indicated time followed by Western blot analysis of indicated proteins. Representative immunoblot images are shown. **(D)** Control RAW264.7 cells or DGAT1-KO RAW264.7 cells were stimulated with HKCA (MOI=3) for 4h followed by immunofluorescence staining of LD with BODIPY and nucleus was stained by DAPI (Scale bar: 20μm), and LD-BODIPY MFI was measured by flow cytometry (Right). Error bars represent SEM of three biological replicates. **(E)** Control THP1 cells or DGAT1-KO THP1 cells were stimulated with HKCA (MOI=3) for 4h followed by immunofluorescence staining of LD with BODIPY and nucleus was stained by DAPI (Scale bar: 20μm), and LD-BODIPY MFI was measured by flow cytometry (Right). Error bars represent SEM of three biological replicates. **(F)** Control THP1 cells or HILPDA-KO THP1 cells were stimulated with HKCA (MOI=3) for 4h followed by immunofluorescence staining of LD with Nile-Red and nucleus was stained by DAPI (Scale bar: 40μm) and LD-Nile-Red MFI was measured by flow cytometry (Right). Error bars represent SEM of three biological replicates. **(G)** *Hilpda*^fl/fl^ or *Hilpda*^fl/fl^ Lyz2-Cre BMDMs cells were stimulated with HKCA (MOI= 3) for 4h followed by immunofluorescence staining of LD with Nile-Red and nucleus was stained by DAPI (Scale bar: 40μm) and LD-Nile-Red MFI was measured by flow cytometry (Right). Error bars represent SEM of four biological replicates. Two-tailed unpaired Student’s t test (D, E, F and G) and one-way ANOVA for (B).

We then performed qPCR analysis to consolidate the RNA-seq data. Upon stimulation with HKCA, D-zymosan or Curdlan, *Dgat1* expression was significantly increased (Figure 2B) and other genes involved in de novo synthesis of triglycerides were also significantly up-regulated, such as *Fasn*, *Agpat4* and *Acsl1* (Figure S2B). Furthermore, the *Atgl* expression induced by curdlan, D-zymosan, or HKCA was significantly reduced and *Hilpda* transcription was significantly increased conversely (Figure 2B). Western blot analysis confirmed that the protein level of DGAT1, HILPDA was increased and ATGL was reduced upon HKCA, D-zymosan or Curdlan induction (Figure 2C). Meanwhile, we observed that hypoxia inducible factor *Hif1α* was also significantly up-regulated upon fungal induction (Figure 2A, S2A and S2B), which may promote the expression of HILPDA and lead to LDs accumulation. Further, the structural proteins involved in the assembly of lipid droplets, PLIN2 and PLIN3 were also induced upon fungal stimulation (Figure 2A, S2B and S2C). Together, these data confirmed that fungal infection induces the expression of genes involved in lipid biosynthesis and inhibited the expression of genes related to lipolysis, which may result in intracellular LDs accumulation in macrophages.

To further verify these conclusions, we first utilized the inhibitors of DGAT1, A922500 or T863, to block triglycerides biosynthesis.^27^ Consistent with the function of DGAT1 in triglycerides biosynthesis, intracellular LD formation in A922500- or T863-pretreated THP1 cells was markedly reduced upon stimulation with HKCA (Figure S2E). Next, we generated DGAT1-deficient RAW264.7 macrophages using the CRISPR-Cas9 technique (Figure S2D). We found that LD formation induced by HKCA or without HKCA stimulation was dramatically decreased in DGAT1 KO RAW 264.7 cells (Figure 2D). We also established DGAT1-deficient THP1 cells using the CRISPR-Cas9 technique (Figure S2D). Similarity to the data in RAW264.7 macrophages, DGAT1 deficiency greatly decreased LD formation induced by HKCA or without HKCA stimulation in THP1 cells (Figure 2E). We further exploited ATGL inhibitor Atglistatin to inhibit the hydrolase activity of ATGL to reduce LD lipolysis. Intracellular LD formation was substantially increased in Atglistatin-pretreated THP1 cells after HKCA stimulation, compared to that in control-treated THP1 cells (Figure S2F). As mentioned above, HILPDA is a natural short peptide which can inhibit ATGL activity and facilitate LD accumulation.^28, 29^ We then generated HILPDA-deficient THP1 cells using the CRISPR-Cas9 technique as well (Figure S2D). Consistently, we found that HILPDA-KO THP1 cells displayed markedly decreased formation of LD indicated staining with Nile-Red after HKCA stimulation (Figure 2F). To further investigate the function of HILPDA in the regulation of LD formation, the *Hilpda*^fl/fl^ mice were crossed with Lyz2-Cre transgenic mice to produce myeloid-specific HILPDA knockout mice. BMDMs were prepared from *Hilpda*^fl/fl^ Lyz2-Cre and *Hilpda*^fl/fl^ mice followed stimulation with HKCA. Consistent with the observation from sgRNA knockout of HILPDA in THP1 cells, stimulation of *Hilpda*^fl/fl^ Lyz2-Cre BMDMs with HKCA led to a significant decrease in the formation of LD compared with that in *Hilpda*^fl/fl^ BMDMs (Figure 2G). Collectively, these data suggested that the promotion of triglyceride synthesis and repression of triglyceride lipolysis contribute to LD accumulation in macrophages during fungal infection.

### LD accumulation restricts phagosome formation upon fungal infection

In an effort to understand the specific function(s) of LD in anti-fungal immunity, we first examined whether LD formation have any effect on phagocytosis. We used HKCA containing GFP expression vector to stimulate macrophages or THP1 cells followed by flow cytometry to measure percentage of GFP^+^ cells as indication of phagocytosis efficiency. We found treatment with A922500 or T863 and Atglistatin, DGAT1 inhibitors and ATGL inhibitor respectively, or knockout of DGAT1 and HILPDA in macrophages or THP1 cells didn’t not affect phagocytosis efficiency (Figure S3A). To directly investigate the function of LD in phagocytosis, we next used oleic acid (OA) to treat THP1 cells or BMDMs to increase LD accumulation. Consistently, LD accumulation labeled with BODIPY was dramatically increased after treatment of OA (Figure S3B and S3C). While, we did not detect any differences in phagocytic efficiency regardless of OA concentration (Figure S3D and S3E). These data suggested that phagocytosis efficiency is not affected by LD accumulation. So, we next explored whether LD accumulation affected phagocytic index or phagosomes number of single cell. Surprisingly, using control RAW264.7 cells or DGAT1-KO RAW264.7 cells to engulf ample HKCA-GFP (MOI=10), we found the phagosomes number of single cell was significantly increased in DGAT-1 deficient RAW264.7 cells, which contained lower LD level (Figure 3A). Furthermore, we employed FITC and Fluor-647 to dual label HKCA and analyzed the cell subpopulations containing varying numbers of phagosomes based on the methods outlined in “Dendritic Cell Protocols” by Shalin H. Naik.^30^ Flow cytometry results indicated significant increases in both population a (cells with a small number of phagosomes) and population b (cells with quite a few phagosomes) in DGAT-1 deficient RAW264.7 cells, after engulfing FITC and Fluor-647 labeled HKCA for 2h or 4h. While the population c (cells with moderate number of phagosomes) had an upward trend but no significant difference between DGAT-1 deficient RAW264.7 cells and control RAW264.7 cells (Figure 3B). Moreover, we isolated BMDMs from *Hilpda*^fl/fl^ Lyz2-Cre and *Hilpda*^fl/fl^ mice followed pretreatment with low-dose D-zymosan (10μg/mL) for 4h to induce LD production and then engulfment with sufficient HKCA-GFP. Consistent with the results from sgRNA knockout of DGAT1 in RAW264.7 cells, the phagosomes number of single cell was also significantly increased in *Hilpda*^fl/fl^ Lyz2-Cre BMDM cells, which induced a few LD production (Figure 3C). Besides, after engulfed FITC and Fluor-647 labeled HKCA for 1h or 2h, the population c and population d (cells with a large numbers of phagosomes) of *Hilpda*^fl/fl^ Lyz2-Cre BMDM cells were enhanced obviously, compared to *Hilpda*^fl/fl^ BMDMs (Figure 3D). These data suggested that LD may regulate the number of phagosomes during fungal infection.

**Figure 3.**
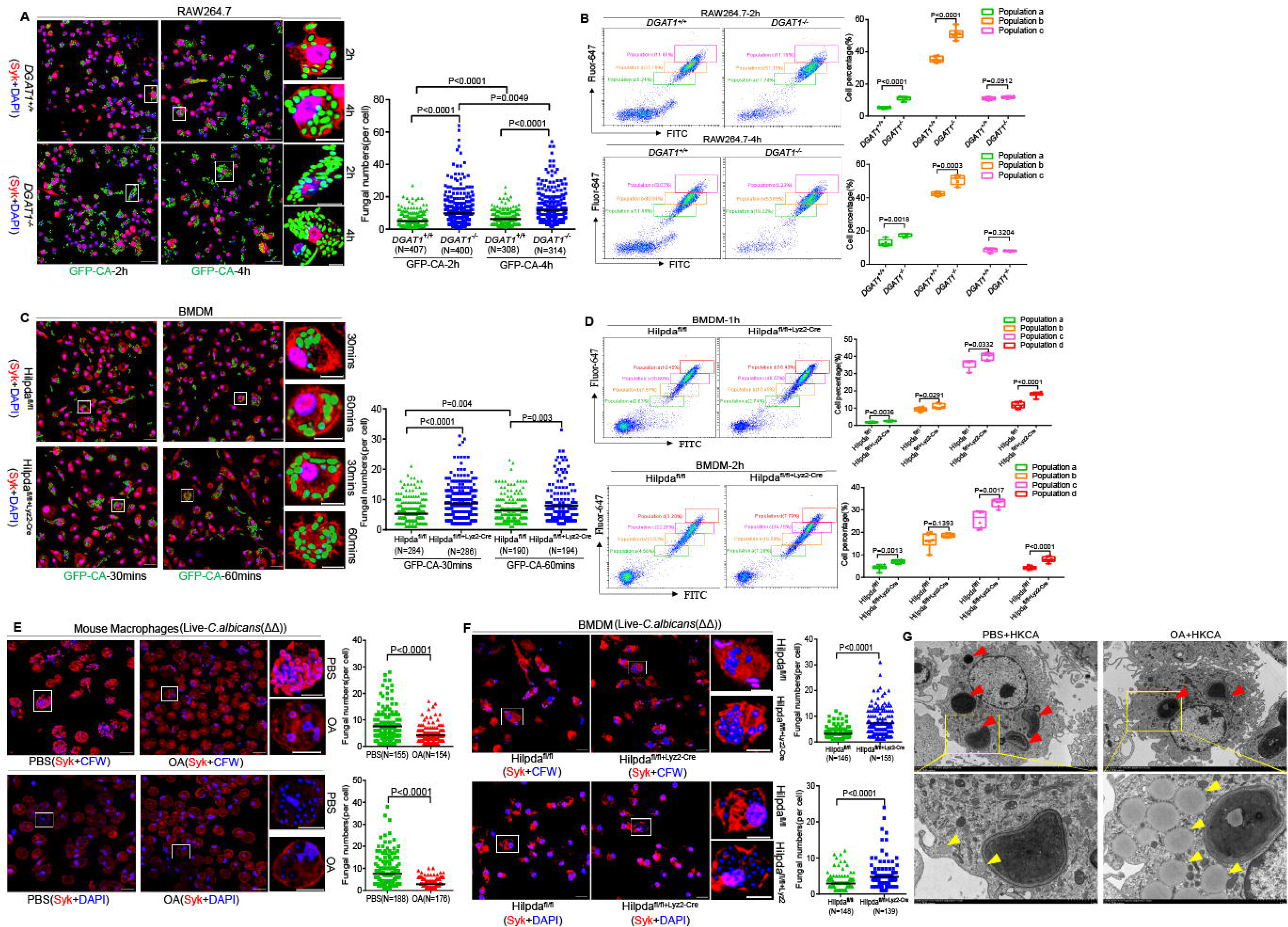
LD accumulation attenuated the number of phagosomes formation. **(A)** Control RAW264.7 cells or DGAT1-KO RAW264.7 cells engulfed HKCA-GFP (MOI=10) for the indicated times followed by immunofluorescence staining of indicated proteins (Scale bar: 40μm and 10μm for representative single cell images). Nuclei were stained with DAPI and Syk staining indicated cell position. The number of phagosomes containing HKCA-GFP was counted in single cell (lower). N represented cell number. **(B)** Control RAW264.7 cells or DGAT1-KO RAW264.7 cells engulfed FITC and Fluor-647 double labeled HKCA (MOI=10) for 2h or 4 h followed by flow cytometry. A detailed analysis of the phagocytic population in the same plot makes evident the presence of different cell populations according to the number of phagosomes internalized by an individual cell (population a: cells with a small number of phagosomes; population b: cells with quite a few phagosomes; population c: cells with moderate number of phagosomes; population d: cells with a large number of phagosomes). Error bars represent SEM of six biological replicates. **(C)** *Hilpda*^fl/fl^ or *Hilpda*^fl/fl^ Lyz2-Cre BMDMs cells were pretreated with low-dose D-zymosan (10μg/mL) for 4h to induce LD production and then engulfed HKCA-GFP (MOI=10) for the indicated times followed by immunofluorescence staining of indicated proteins (Scale bar: 40μm and 10μm for representative single cell images). Nuclei were stained with DAPI and Syk staining indicated cell position. The number of phagosomes containing HKCA-GFP was counted in single cell (lower). N represented cell number. **(D)** *Hilpda*^fl/fl^ or *Hilpda*^fl/fl^ Lyz2-Cre BMDMs cells were pretreated with low-dose D-zymosan (10μg/mL) for 4h to induce LD production and then engulfed FITC and Fluor-647 double labeled HKCA (MOI=10) for 1h or 2h followed by flow cytometry. A detailed analysis of the phagocytic population in the same plot makes evident the presence of different cell populations according to the number of phagosomes internalized by an individual cell (population a: cells with a small number of phagosomes; population b: cells with quite a few phagosomes; population c: cells with moderate number of phagosomes; population d: cells with a large number of phagosomes). Error bars represent SEM of six biological replicates. **(E)** Mouse macrophages were pretreated with OA (4.5μg/mL) for 24h, then engulfed live hyphae-deficient (cph1Δ/Δ, efg1Δ/Δ) *C. albicans* (MOI=10) for 24h followed by immunofluorescence staining of indicated proteins (Scale bar: 20μm and 10μm for representative single cell images). The number of phagosomes formation was counted in single cell (Right). Syk indicated cell position, engulfed fungi were stained with CFW (upper) or DAPI (down) and N represented cell number. **(F)** *Hilpda*^fl/fl^ or *Hilpda*^fl/fl^ Lyz2-Cre BMDMs cells were pretreated with low-dose D-zymosan (10μg/mL) for 4h to induce LD production, then engulfed live hyphae-deficient (cph1Δ/Δ, efg1Δ/Δ) *C. albicans* (MOI=10) for 24h followed by immunofluorescence staining of indicated proteins (Scale bar: 20μm and 10μm for representative single cell images). The number of phagosomes formation was counted in single cell (Right). Syk indicated cell position, engulfed fungi were stained with CFW (upper) or DAPI (down) and N represented cell number. **(G)** BMDMs cells were pretreated with OA (4.5μg/mL) for 24h, then stimulated with HKCA (MOI=5) for 4h followed by electron microscopy images. LDs are indicated by yellow arrowheads and phagosomes are indicated by red arrowheads. Two-tailed unpaired Student’s t test (A-F).

To further confirm these phenomena, we then counted the phagosome number in OA-pretreated THP1 cells followed stimulation with HKCA-GFP. We found the phagosome number in OA-treated THP1 cells was dramatically decreased compared to control group (Figure S4A), and the population c and population d of OA-treated BMDM cells were significantly declined (Figure S4D). Similarly, treatment with ATGL inhibitor Atglistatin to increase intracellular LDs also reduced the phagosome number in THP1 cells stimulated with HKCA-GFP compared to control group (Figure S4B), and the population c and population d of Atglistatin-treated BMDM cells were also significantly reduced (Figure S4E). In contrast, THP1 cells pretreated with DGAT1 inhibitors A922500 or T863 to decrease intracellular LDs showed increased phagosome number compared to control group (Figure S4C), and the population c and population d of A922500 or T863-treated BMDM cells were significantly expanded (Figure S4F).

To further explore this hypothesis, we chose live hyphae-deficient (cph1Δ/Δ, efg1Δ/Δ) *C. albicans* to perform these experiments as wild-type *C. albicans* could form hyphae and escape from macrophages.^18,19^ The phagosomes were stained by two methods using DAPI-labeled fungal nucleus and calcofluor white (CFW)-labeled fungal cell wall (Figure 3E and 3F). We found LD accumulation induced by OA in mouse macrophages could markedly reduce the number of phagosomes after live hyphae-deficient (cph1Δ/Δ, efg1Δ/Δ) *C. albicans* infection for 24h (Figure 3E). Conversely, the number of phagosomes was dramatically raised in *Hilpda*^fl/fl^ Lyz2-Cre BMDM cells after live hyphae-deficient (cph1Δ/Δ, efg1Δ/Δ) *C. albicans* infection for 24h, compared to *Hilpda*^fl/fl^ BMDM cells (Figure 3F). Furthermore, the images of transmission electron microscopy showed that the LDs induced by OA could reduce phagosomes formation in BMDMs stimulated with HKCA for 30 minutes (Figure 3G). Taken together, these data suggested that LD accumulation attenuates the quantity of phagosomes after fungal infection.

Next, we explored whether LD is involved in antifungal signaling activation. We found induction of LD accumulation by OA attenuated NF-κB and MAPK signaling activation and proinflammatory cytokines expression upon Dectin ligand stimulation (data not show).

### LD accumulation and phagosome formation competes in utilizing intracellular endoplasmic reticulum membrane components

In the process of phagocytosis of fungi, the Dectin receptors on the cell membrane of macrophages combines with the fungal cell wall components, and gradually sinks into the cell, then the membrane components in the cell (mainly the endoplasmic reticulum membrane) supplement and fuse with the phagosome membrane.^12^ Our recent study also demonstrated that endoplasmic reticulum membrane may function as an important phagosomal membrane source and the endoplasmic reticulum protein STING translocates to the phagosomes to negatively regulate anti-fungal immunity.^17^ The formation of LD occurs in the ER, and the triacylglycerides were wrapped in a single layer membrane. However, this single layer membrane was also provided by the ER.^20^ According to these reports, we hypothesized that during the formation process of phagosomes and LDs, they may compete for membrane source from the ER.

To test this hypothesis, we first examined whether controlling LD level could affect the number of phagosomes formation. We pretreated THP1 cells with OA or Atglistatin to increase intracellular LDs followed with HKCA stimulation. Consistently, the surface of accumulated LDs was colocalized with endoplasmic reticulum membrane labeled by Calnexin. Notably, the number of phagosomes formation was obviously decreased in OA- or Atglistatin-treated cells (Figure 4A and 4B). We further pretreated THP1 cells with A922500 or T863 to decrease intracellular LDs. We showed that massive endoplasmic reticulum membrane components in the cell were not colocalized with LDs, but with more phagosomes (Figure 4C). Similarly, massive endoplasmic reticulum membrane components in the cell were not occupied by LDs and they were used to form more phagosomes in DGAT-1 deficient RAW264.7 cells and THP1 cells, (Figure 4D and 4E). Moreover, in *Hilpda*^fl/fl^ Lyz2-Cre BMDMs, since LDs was not induced much more by low-dose D-zymosan, HKCA infection made the endoplasmic reticulum membrane components to be used for forming more phagosomes, compared to the BMDMs from *Hilpda*^fl/fl^ mice (Figure 4F). These data suggest that the preferential formation of lipid droplets might control phagosomes formation.

**Figure 4.**
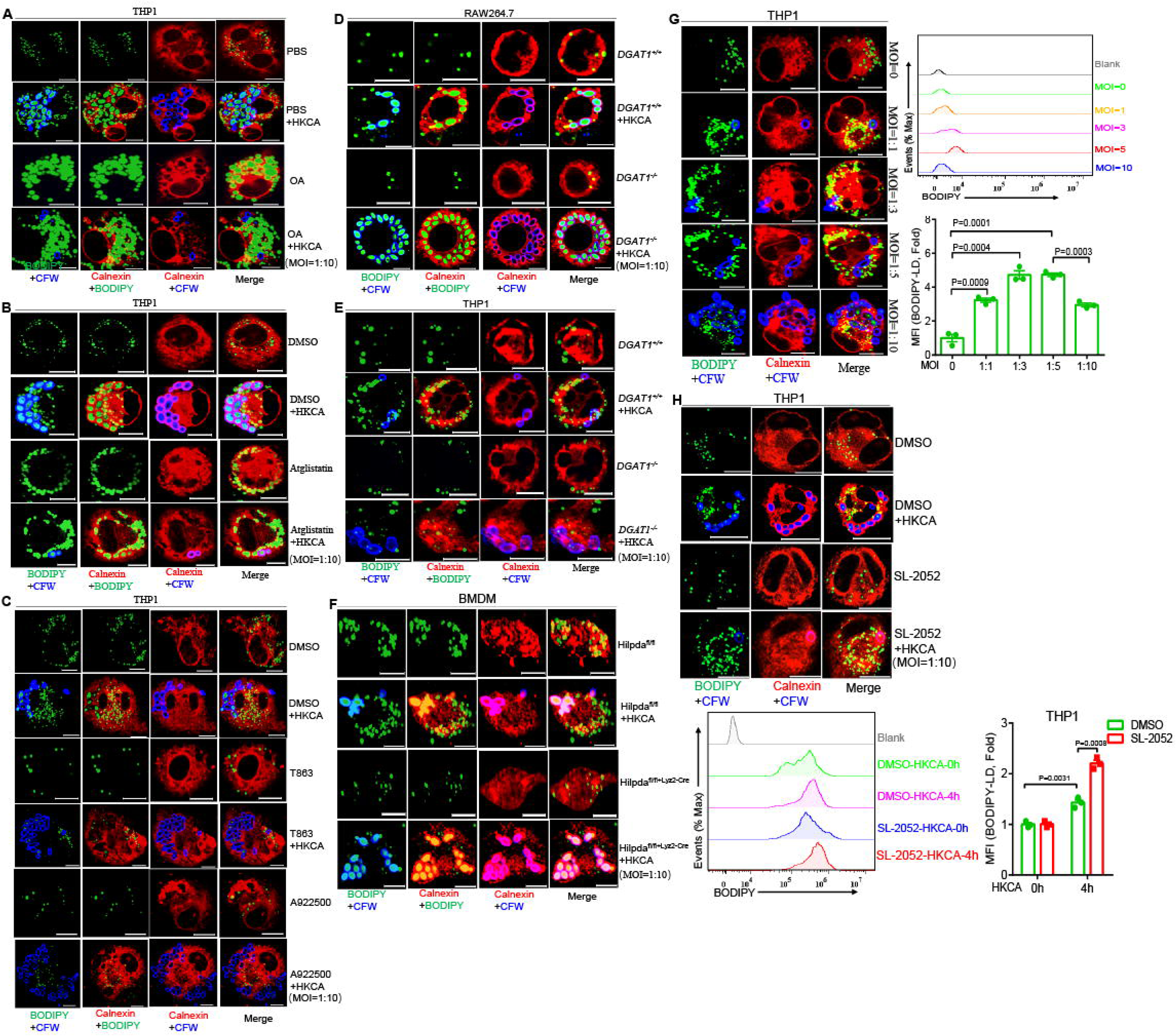
LD accumulation and phagosomes formation competed in utilizing cell membrane components. **(A)** THP1 cells were pretreated with or without OA (4.5μg/mL) for 24h, then engulfed HKCA (MOI=10) for 4h followed by immunofluorescence staining of indicated proteins (Scale bar: 10μm). Calnexin indicated endoplasmic reticulum, CFW stained fungal cell wall and LD was stained by BODIPY. **(B)** THP1 cells were pretreated with or without Atglistatin (10μM) for 24h, then engulfed HKCA (MOI=10) for 4h followed by immunofluorescence staining of indicated proteins (Scale bar: 10μm). Calnexin indicated endoplasmic reticulum, CFW stained fungal cell wall and LD was stained by BODIPY. **(C)** THP1 cells were pretreated with or without A922500 (10μM) or T863 (10μM) for 24h, then engulfed HKCA (MOI=10) for 4h followed by immunofluorescence staining of indicated proteins (Scale bar: 10μm). Calnexin indicated endoplasmic reticulum, CFW stained fungal cell wall and LD was stained by BODIPY. **(D)** Control RAW264.7 cells or DGAT1-KO RAW264.7 cells engulfed HKCA (MOI=10) for 4h followed by immunofluorescence staining of indicated proteins. Calnexin indicated endoplasmic reticulum, CFW stained fungal cell wall and LD was stained by BODIPY. **(E)** Control THP1 cells or DGAT1-KO THP1 cells engulfed HKCA (MOI=10) for 4h followed by immunofluorescence staining of indicated proteins. Calnexin indicated endoplasmic reticulum, CFW stained fungal cell wall and LD was stained by BODIPY. **(F)** *Hilpda*^fl/fl^ or *Hilpda*^fl/fl^ Lyz2-Cre BMDMs cells were pretreated with low-dose D-zymosan (10μg/mL) for 4h to induce LD production and then were stimulated with HKCA (MOI = 10) for 30 minutes followed by immunofluorescence staining of indicated proteins (Scale bar: 10μm). Calnexin indicated endoplasmic reticulum, CFW stained fungal cell wall and LD was stained by BODIPY. **(G)** THP1 cells were stimulated with different infection ratio of HKCA followed by immunofluorescence staining of indicated proteins (Scale bar: 10μm) and flow cytometry to detect the LD-BODIPY MFI. Calnexin indicated endoplasmic reticulum, CFW stained fungal cell wall and LD was stained by BODIPY. Error bars represent SEM of three biological replicates. **(H)** THP1 cells were pretreated with or without SL-2052 (10μM) for 1h, then stimulated with HKCA (MOI=10) for 4h followed by immunofluorescence staining of indicated proteins (Scale bar: 10μm), and flow cytometry to detect the LD-BODIPY MFI Calnexin indicated endoplasmic reticulum, CFW stained fungal cell wall and LD was stained by BODIPY. Error bars represent SEM of three biological replicates. Two-tailed unpaired Student’s t test (H) and one-way ANOVA for (G).

Next, we explored whether phagosomes formation could affect LDs accumulation. THP1 cells were engulfed with different multiplicity of infection (MOI) of HKCA, then intracellular LDs level was detected by confocal microscopy and flow cytometry. Consistently, more phagosomes were formed in macrophages with the increase of MOI HKCA infection. LDs level were also gradually increased as HKCA MOI increased. Interestingly, the maximum level of LDs was at 3 MOI of HKCA, and intracellular LD level was not further increased at 5 MOI infection. Notably, intracellular LD level was significantly decreased when maximum phagosomes were formed with 10 MOI HKCA (Figure 4G). These results were confirmed by repeating experiments on mouse macrophages and showed similar trends (Figure S5A and S5B). Phosphatidylinositol 3-kinase (PI3K)/Akt signaling pathway activation plays important role in phagocytosis.^12^ To further demonstrate phagosome formation affects LD accumulation, we used PI3 kinase inhibitors, which can dampen phagocytosis. We chose three inhibitors, Ly294002, wortmannin (SL-2052) and XL-147. Phagocytosis assay showed that the inhibitory effect of SL-2052 was most efficient (Figure S5C). Moreover, SL-2052 pretreatment did not affect the genes transcription and proteins expression involved in LD formation during HKCA stimulation, such as HILPDA, DGAT-1, ATGL and PLIN3 (Figure S5D and S5E). With the decrease of phagosome formation, we found intracellular LD level in THP1 cells pretreated with SL-2052 was markedly enhanced after HKCA infection (Figure 4H). These data indicate that the formation of phagosomes could attenuate LD formation as well. Taken together, we found that competitive consumption of intracellular endoplasmic reticulum membrane components by LD might contribute to restrict phagosomes formation in macrophages during fungal infection.

### LD formation restricts the quantity of phagosomes via altering RAC1 GTPase activity

PI3K/Akt signaling plays important role in phagocytosis.^12^ So, we first examined whether LD accumulation could affect the PI3K/Akt activation during fungal infection. Pretreatment with OA to induce intracellular LD accumulation did not impair PI3K/Akt signaling as showed by P-Akt (T-308) and P-Akt (S-473) level in THP1 cells, RAW264.7 macrophages or BMDMs cell during HKCA stimulation (Figure S6A-S6C). To further understand the specific molecular mechanism of LD regulating phagosomes formation, we isolated and purified intracellular LDs followed by mass spectrometry to identify LD surface proteins. Using this method, we identified Rho family small GTP enzymes RAC1/2 were assembled to LD surface after OA induction or HKCA stimulation (Figure 5A). Confocal microscopy also showed that RAC1 could locate on LD surface (Figure S6D) and colocalize with LD structural protein PLIN2 (Figure S6E). Western blot analysis with the purified LDs showed that RAC1 was recruited to LD upon both OA induction and HKCA stimulation (Figure S6F and S6G). When macrophages engulf fungi, the mycelium will sink inward with the cell membrane, which requires the aggregation of microfilaments to drive the outward extension and encapsulation of cell membrane. Importantly, the aggregation of microfilaments requires the activation of Rho family small GTPases, including RAC1 and CDC42.^14^ In addition, a recent study also identified that RAC1 and CDC42 were clustered into LDs during LPS stimulation.^26^ According to these observations, we speculated that LD may affect the formation of phagosomes by changing the localization of RAC1 from phagosomes to LD and regulating its activity during phagocytosis. To prove this hypothesis, we first evaluated RAC GTPase activity and found that RAC1 activation was greatly increased in THP1 cells after HKCA stimulation but was abolished in THP1 cells treated with OA (Figure 5B). In contrast, RAC1 activation was markedly boosted in THP1 cells pretreated with A922500 or T863 after HKCA stimulation (Figure 5C and 5D). Besides, the RAC1 GTPase activity was also enhanced in DGAT-1 deficient RAW264.7 cells after HKCA stimulation, compared to control RAW264.7 cells (Figure 5E). These data indicated that LD accumulation weakens RAC1 activation during fungal infection. Further, confocal microscopy showed that RAC1 was recruited to cell membrane at early time stage (0.5h and 1h) and facilitated the subsequent phagocytosis in control THP1 cells upon fungi infection (Figure 5F). This result was consistent with a previous report that HKCA or α-Mannan stimulation triggered RAC1 translocation to cell membrane.^31^ However, OA treatment “locked” RAC1 on LD surface and was unable to translocate to the cell membrane in THP1 cells regardless of HKCA stimulation (Figure 5F). Subcellular fractionation experiments also displayed that RAC1 membrane translocation was obviously suppressed during HKCA infection in THP1 cell treated with OA (Figure 5G). In addition, the RAC1 membrane translocation was risen in DGAT-1 deficient RAW264.7 cells after HKCA stimulation (Figure 5H). HKCA treatment increased the interaction between RAC1 and LD structural protein PLIN3, indicating PLIN3 may recruit RAC1 to LD surface (Figure 5I and 5J). Together, our results suggested that LD accumulation could limit phagosomes formation through regulating RAC1 translocation and its GTPase activity.

**Figure 5.**
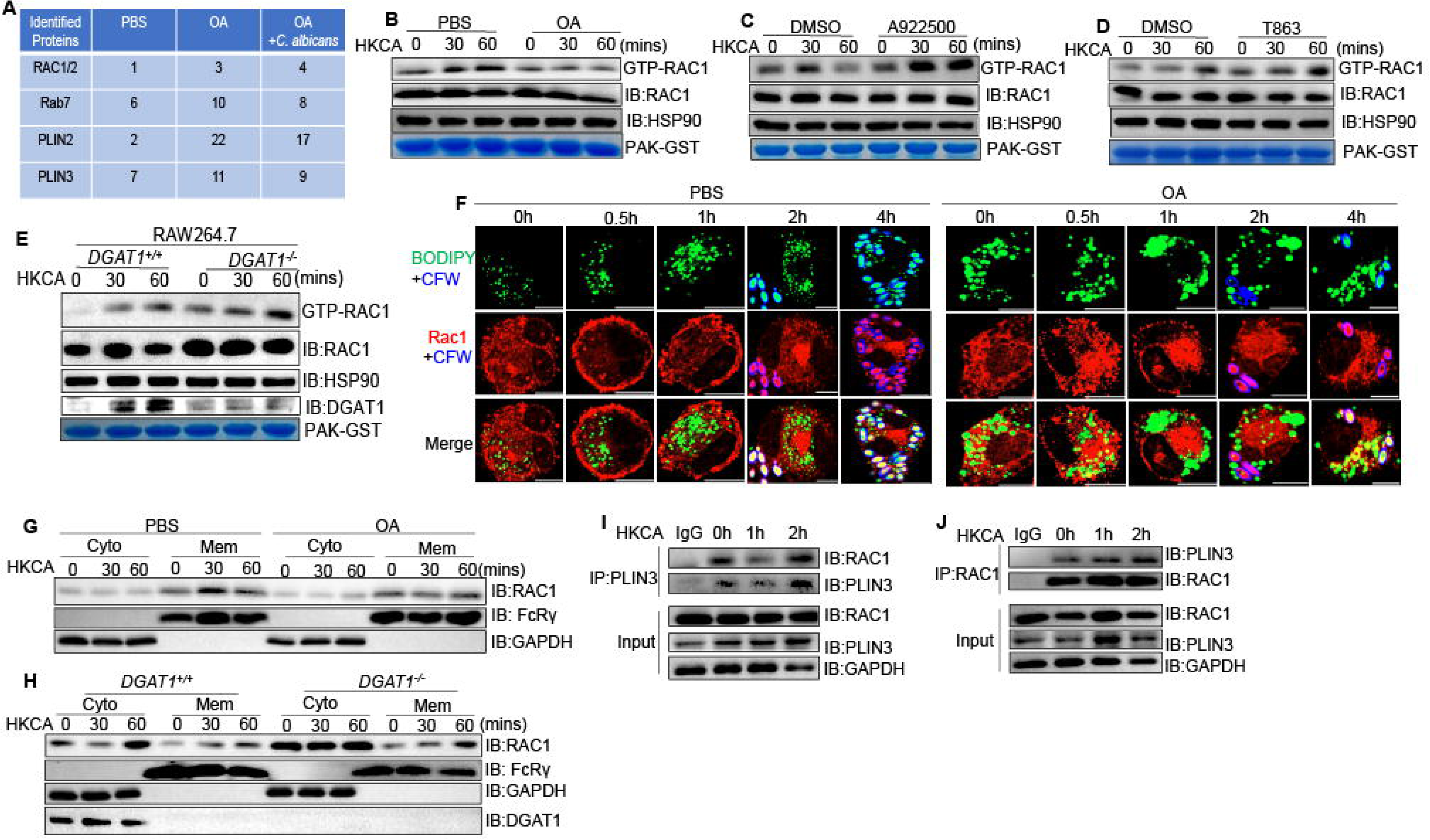
LD formation restricts the quantity of phagosomes via altering RAC GTPase activity. **(A)** THP1 cells were pretreated with or without OA (4.5μg/mL) for 24h, then engulfed HKCA (MOI=10) for 4h followed by immunofluorescence staining of indicated proteins (Scale bar: 10μm). Calnexin indicated endoplasmic reticulum, CFW stained fungal cell wall and LD was stained by BODIPY. **(B)** THP1 cells were pretreated with or without Atglistatin (10μM) for 24h, then engulfed HKCA (MOI=10) for 4h followed by immunofluorescence staining of indicated proteins (Scale bar: 10μm). Calnexin indicated endoplasmic reticulum, CFW stained fungal cell wall and LD was stained by BODIPY. **(C)** THP1 cells were pretreated with or without A922500 (10μM) or T863 (10μM) for 24h, then engulfed HKCA (MOI=10) for 4h followed by immunofluorescence staining of indicated proteins (Scale bar: 10μm). Calnexin indicated endoplasmic reticulum, CFW stained fungal cell wall and LD was stained by BODIPY. **(D)** Control RAW264.7 cells or DGAT1-KO RAW264.7 cells engulfed HKCA (MOI=10) for 4h followed by immunofluorescence staining of indicated proteins. Calnexin indicated endoplasmic reticulum, CFW stained fungal cell wall and LD was stained by BODIPY. **(E)** Control THP1 cells or DGAT1-KO THP1 cells engulfed HKCA (MOI=10) for 4h followed by immunofluorescence staining of indicated proteins. Calnexin indicated endoplasmic reticulum, CFW stained fungal cell wall and LD was stained by BODIPY. **(F)** *Hilpda*^fl/fl^ or *Hilpda*^fl/fl^ Lyz2-Cre BMDMs cells were pretreated with low-dose D-zymosan (10μg/mL) for 4h to induce LD production and then were stimulated with HKCA (MOI = 10) for 30 minutes followed by immunofluorescence staining of indicated proteins (Scale bar: 10μm). Calnexin indicated endoplasmic reticulum, CFW stained fungal cell wall and LD was stained by BODIPY. **(G)** THP1 cells were stimulated with different infection ratio of HKCA followed by immunofluorescence staining of indicated proteins (Scale bar: 10μm) and flow cytometry to detect the LD-BODIPY MFI. Calnexin indicated endoplasmic reticulum, CFW stained fungal cell wall and LD was stained by BODIPY. Error bars represent SEM of three biological replicates. **(H)** THP1 cells were pretreated with or without SL-2052 (10μM) for 1h, then stimulated with HKCA (MOI=10) for 4h followed by immunofluorescence staining of indicated proteins (Scale bar: 10μm), and flow cytometry to detect the LD-BODIPY MFI Calnexin indicated endoplasmic reticulum, CFW stained fungal cell wall and LD was stained by BODIPY. Error bars represent SEM of three biological replicates. Two-tailed unpaired Student’s t test (H) and one-way ANOVA for (G).

### LD accumulation protects macrophages from death

After phagocytosis of live *C. albicans*, fungi initially could escape from macrophages through caspase-1-dependent pyroptosis, followed by massive macrophage death.^18^ During the process, GSDMD activation also facilitates escape of *C. albicans* from macrophages by punching holes on the cell membrane. Meanwhile, *C. albicans* deployed hyphae form and secreted candidalysin, a pore-forming toxin to physically damage cell membrane and escape (19). Besides, one recent research found that the hscA deletion mutant (ΔhscA) of *Aspergillus fumigatus* attenuated its infection capacity into lung epithelial cells and the number of ΔhscA conidia within per A549 cells was reduced compared with WT conidia. Subsequently, the ΔhscA conidia were easier to be cleared by Rab7-mediated phagolysosome maturation.^32^ So, we hypothesized that controlled phagosomes formation by LD accumulation may protect macrophages from death to increase the elimination efficiency of fungi infection. First, we infected BMDM cells with different MOI of live wild type *C. albicans* and found that the death of macrophages was gradually increased with the increase of MOI *C. albicans* infection (Figure 6A). Additionally, consistent with other reports, the formation of *C. albicans* hyphae caused higher macrophage death compared to hyphae-deficient (cph1Δ/Δ, efg1Δ/Δ) *C. albicans* (Figure 6B, 6C and S7A). Surprisingly, OA treatment could significantly reduce LDH release from THP1 cell infected by both wild-type *C. albicans* and hyphae-deficient (cph1Δ/Δ, efg1Δ/Δ) *C. albicans* (Figure 6B). The cell death of THP1 cells or BMDMs also decreased by OA treatment as stained with PI dye or fixable viability dye, respectively (Figure 6C and 6D). Besides, pretreatment with Atglistatin also decreased LDH release from THP1 cells and reduced cell death after fungal infection (Figure S7B-S7D). In contrast, knocked out of DGAT-1 in RAW264.7 cells or pretreatment with A922500 or T863 of THP1 cells increased the LDH release, and more cell death was showed as stained with PI dye (Figure 6E, 6F and S7E-S7G). Consistent with the results from sgRNA knockout of DGAT1 in RAW264.7 cells, infection of *Hilpda*^fl/fl^ Lyz2-Cre BMDMs with live *C. albicans* led to a significant increase of LDH release and cell death, compared with that in *Hilpda*^fl/fl^ BMDMs (Figure 6G and 6H). Together, these data suggested that LD play a crucial role on protecting macrophages from death.

**Figure 6.**
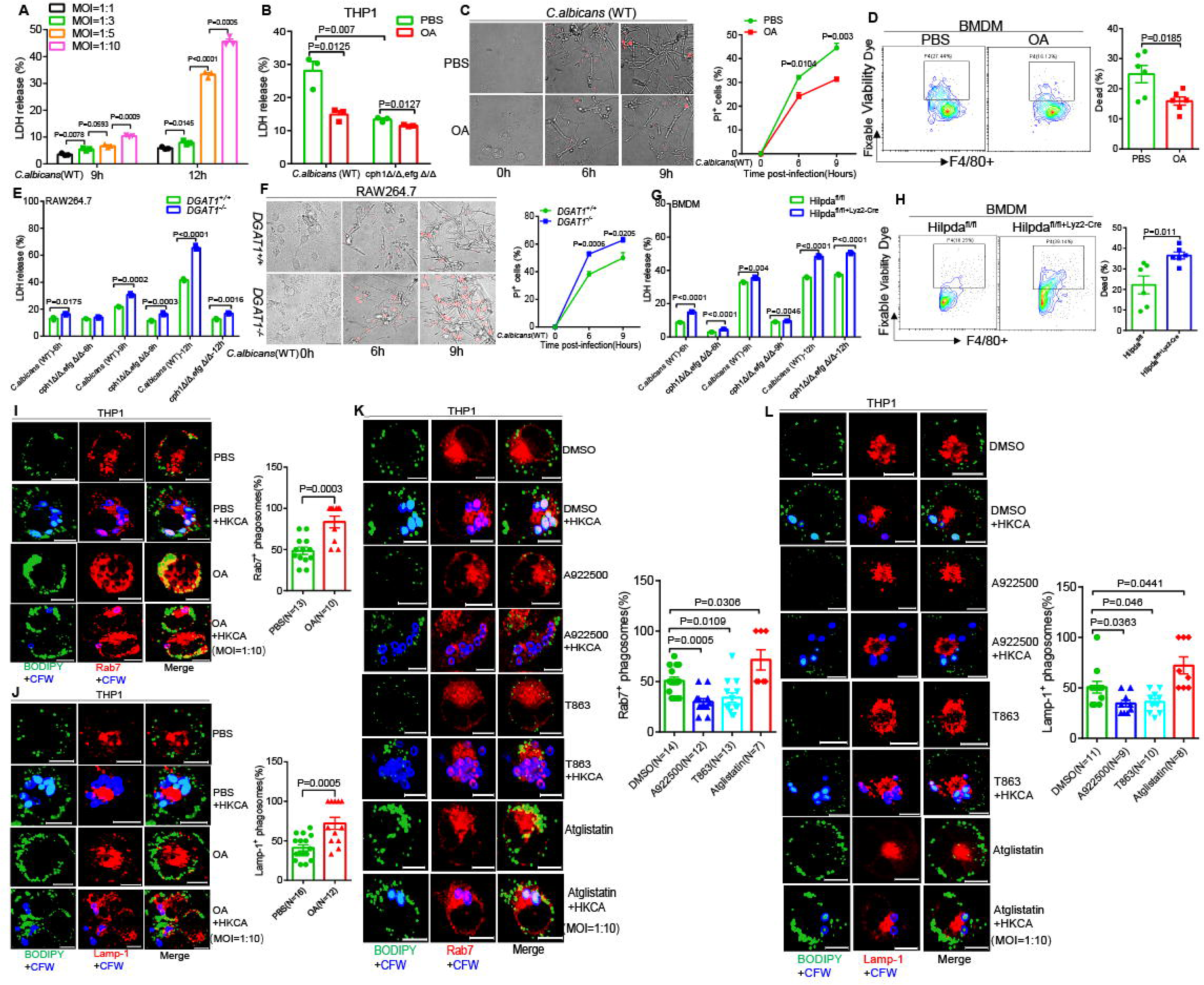
Controlled phagosomes formation by LD accumulation protected macrophages from death. **(A)** THP1 cells were infected with different infection ratio of live wild-type *C. albicans* for the indicated time followed by a lactate dehydrogenase (LDH) cytotoxicity assay of THP1 cells. Error bars represent SEM of three biological replicates. **(B-C)** THP1 cells were pretreated with or without OA (4.5μg/mL) for 24h, then were infected with live wild-type *C. albicans* or hyphae-deficient (cph1Δ/Δ, efg1Δ/Δ) *C. albicans* (MOI=10) for the indicated time followed by a lactate dehydrogenase (LDH) cytotoxicity assay of THP1 cells, and dying THP1 cells were stained with PI (red) (Scale bar: 20μm). Error bars represent SEM of three biological replicates. **(D)** BMDMs cells were pretreated with or without OA (4.5μg/mL) for 24h, then were infected with live wild-type *C. albicans* for 9h followed by flow cytometry to test cell viability. Error bars represent SEM of six biological replicates. **(E-F)** Control RAW264.7 cells or DGAT1-KO RAW264.7 cells were infected with live wild-type *C. albicans* or hyphae-deficient (cph1Δ/Δ, efg1Δ/Δ) *C. albicans* (MOI=10) for the indicated time followed by a lactate dehydrogenase (LDH) cytotoxicity assay of RAW264.7 cells (E), and dying RAW264.7 cells were stained with PI (red) (Scale bar: 20μm, F). Error bars represent SEM of three biological replicates. **(G)** *Hilpda*^fl/fl^ or *Hilpda*^fl/fl^ Lyz2-Cre BMDMs cells were infected with live wild-type *C. albicans* or hyphae-deficient (cph1Δ/Δ, efg1Δ/Δ) *C. albicans* (MOI=10) for the indicated time followed by a lactate dehydrogenase (LDH) cytotoxicity assay. Error bars represent SEM of three biological replicates. **(H)** *Hilpda*^fl/fl^ or *Hilpda*^fl/fl^ Lyz2-Cre BMDMs cells were infected with live wild-type *C. albicans* for 9h followed by flow cytometry to test cell viability. Error bars represent SEM of six biological replicates. **(I)** THP1 cells were pretreated with or without OA (4.5μg/mL) for 24h, then engulfed HKCA (MOI=10) for 8h followed by immunofluorescence staining of indicated proteins (Scale bar: 10μm). The Rab7^+^ phagosomes were counted in single cell (Right). N represented cell number, CFW stained fungal cell wall and LD was stained by BODIPY. **(J)** THP1 cells were pretreated with or without OA (4.5μg/mL) for 24h, then engulfed HKCA (MOI=10) for 8h followed by immunofluorescence staining of indicated proteins (Scale bar: 10μm). The Lamp-1^+^ phagosomes were counted in single cell (Right). N represented cell number, CFW stained fungal cell wall and LD was stained by BODIPY. **(K)** THP1 cells were pretreated with DMSO, A922500 (10μM), T863 (10μM) or Atglistatin (10μM) for 24h, then engulfed HKCA (MOI=10) for 8h followed by immunofluorescence staining of indicated proteins (Scale bar: 10μm). The Rab7^+^ phagosomes were counted in single cell (Right). N represented cell number, CFW stained fungal cell wall and LD was stained by BODIPY. **(L)** THP1 cells were pretreated with DMSO, A922500 (10μM), T863 (10μM) or Atglistatin (10μM) for 24h, then engulfed HKCA (MOI=10) for 8h followed by immunofluorescence staining of indicated proteins (Scale bar: 10μm). The Lamp-1^+^ phagosomes were counted in single cell (Left). N represented cell number, CFW stained fungal cell wall and LD was stained by BODIPY. Two-tailed unpaired Student’s t test (B, D, E, G, H, I, J, K and L) and one-way ANOVA for (A, C and F).

To explore the potential mechanism underlying LD’s protective effects from cell death, we analyzed the proteome of LD surface proteins again and found that there were many innate immune proteins clustered on LD, such as dermcidin, histone, antibacterial peptide S100-A8 and lysozyme-C etc (Table S8). These effector proteins could directly kill invading fungi and confirmed the concept that LDs are innate immune hubs integrating cell metabolism and host defense.^26^ Except these effector proteins, we also identified Rab7, a protein mediating endoplasmic transport, enrichment in LD surface after OA induction and HKCA stimulation (Figure 5A, S6F and S6G). Since Rab7 plays an essential role in phagosome maturation (32), we analyzed the presence of Rab7 in phagosomes. When THP1 cells were pretreated with OA or Atglistatin, the phagosomes formation was reduced as expected. However, the percentage of Rab7-positive phagosomes was significantly increased compared to control THP1 cells (Figure 6I and 6K). In turn, THP1 cells pretreated with A922500 or T863 were engulfed more fungi, nevertheless the percentage of Rab7-positive phagosomes was significantly decreased (Figure 6K). These data indicate that Rab7 enriched on LD surface may promote phagosome maturation. Rab7 promotes phagosome maturation via directing phagosome to lysosome and forming phagolysosomes.^32^ So, we further examined downstream phagolysosomes formation. As expected, OA or Atglistatin stimulation promoted the percentage of Lamp1-positive phagosomes in THP1 cells, indicative of increased phagolysosomes formation compared to control THP1 cell (Figure 6J and 6L). In turn, the percentage of Lamp1-positive phagosomes was reduced prominently in THP1 cells pretreated with A922500 or T863 (Figure 6L). Taken together, these data suggested that LD accumulation promotes Rab7-mediated phagolysosome maturation.

### Myeloid *Hilpda* deficient mice were susceptible to fungal sepsis

The accumulation of intracellular LD depends on the balance between lipid synthesis and degradation.^20^ Due to the important role of triglycerides in cellular metabolism, lipid biology, and cellular energy supply, knocking out any metabolic enzyme in lipid synthesis or lipid decomposition may affect the normal survival of cells. In addition, our results found that after fungal infection, the synthesis of lipids were enhanced and the lipid degradation pathway was inhibited. HILPDA is an inhibitor of ATGL which mediates intracellular lipolysis. Expression of HILPDA can also be induced by fungal infection (Figure 2A-2C and S2A). We also confirmed the interaction between ATGL and HILPDA through co-immunoprecipitation (Figure S8A and S8B) and identified the N-terminal (1-31aa) of HILPDA domain mediated binding to ATGL by immunofluorescence study (Figure S8C). In order to definitively confirm the physiological function of LD during fungal sepsis, we generated *Hilpda*^fl/fl^ Lyz2-Cre mice with genetic *Hilpda* deletion in myeloid cells. The knockout efficiency of HILPDA was examined by Western blot after HKCA induction (Figure S8D). *In vivo*, *Hilpda*^fl/fl^ Lyz2-Cre mice were more susceptible to *C. albicans*-induced fungal sepsis than *Hilpda*^fl/fl^ control mice, which was shown by the higher mortality rate, faster weight loss, and higher fungal burden of *Hilpda*^fl/fl^ Lyz2-Cre mice compared to *Hilpda*^fl/fl^ mice (Figure 7A-7C). Histopathological analysis and PAS staining also showed the increasement of fungal burden in the kidneys of *Hilpda*^fl/fl^ Lyz2-Cre mice (Figure 7D). Consistent with the function of HILPDA in the regulation of LD levels and modulating the macrophages death *in vitro*, macrophages from kidney of *Hilpda*^fl/fl^ Lyz2-Cre mice were died more than *Hilpda*^fl/fl^ mice counterpart after *C. albicans* infected 5 days (Figure 7E). This result indicated that *Hilpda* deficiency triggering the death of macrophages via decreasing LD levels contributed to the higher mortality and fungal burden of *Hilpda*^fl/fl^ Lyz2-Cre mice during fungal infection.

**Figure 7.**
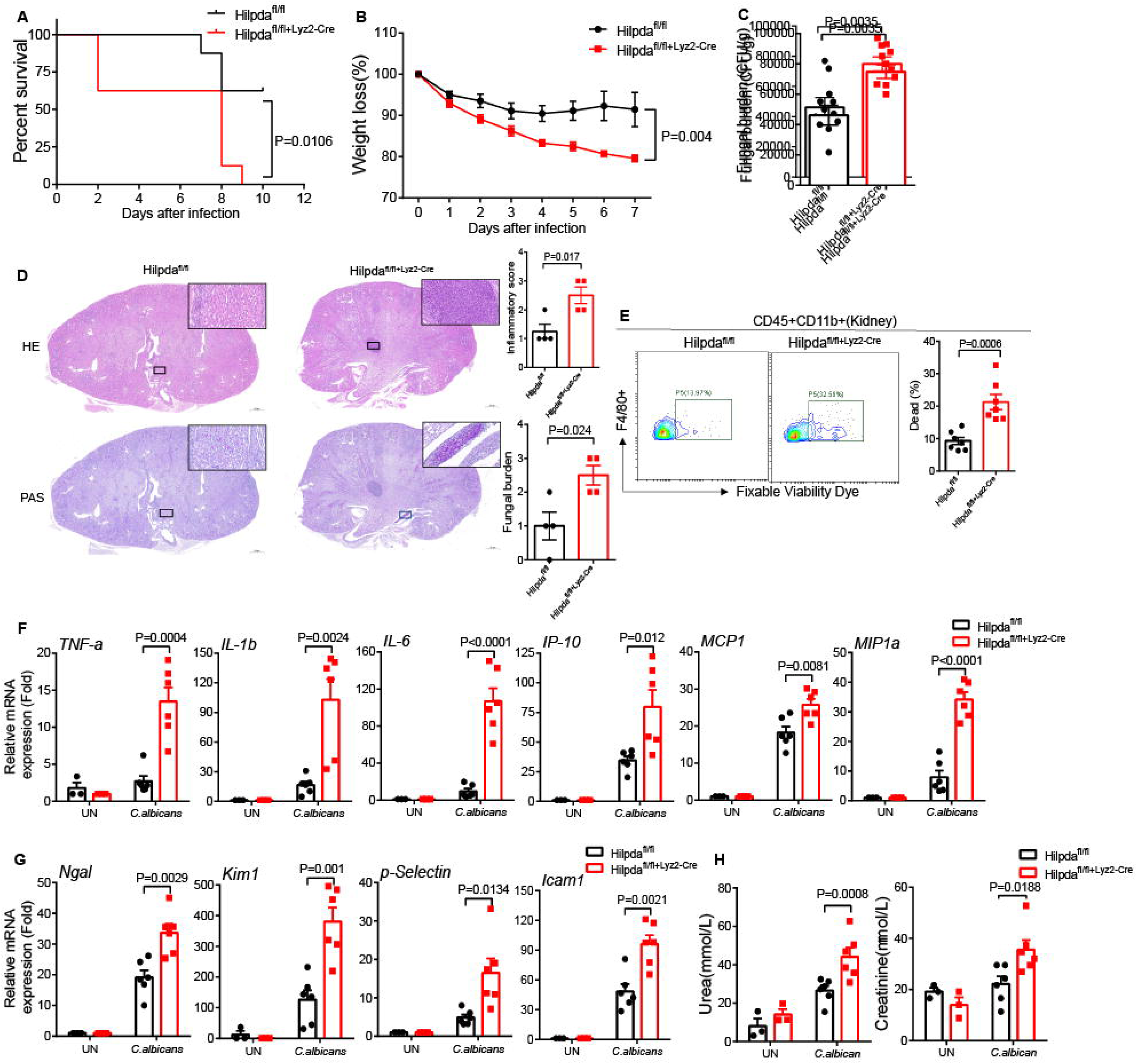
Mice *Hilpda* deletion in macrophage were susceptible to fungal sepsis. **(A-C)** *Hilpda*^fl/fl^ or *Hilpda*^fl/fl^ Lyz2-Cre mice were intravenously injected with live *C. albicans* (4×10^5^cfu/100μL 1×PBS). Survival rate (A), weight loss rate (B), or fungal number (C) in the kidney of mice were shown (n = 8 for A, n = 6 for B, and n = 7 for C). **(D)** *Hilpda*^fl/fl^ or *Hilpda*^fl/fl^ Lyz2-Cre mice were treated as in C, and 5 days after *C. albicans* infection, mice were euthanized, kidneys were fixed, and sections were stained with hematoxylin and eosin (H&E) or periodic-acid Schiff (PAS). Renal inflammation and fungal burden were scored based upon H&E and PAS staining (Right), respectively. Error bars represent SEM of four independent experiments. **(E)** *Hilpda*^fl/fl^ or *Hilpda*^fl/fl^ Lyz2-Cre mice were intravenously injected with live *C. albicans* (4×10^5^ cfu/100 μL 1×PBS), and mice were euthanized at day 5 after infection. The cell viability of macrophages from kidneys were stained for flow cytometry analysis. Error bars represent SEM of seven mice experiments. **(F-G)** *Hilpda*^fl/fl^ or *Hilpda*^fl/fl^ Lyz2-Cre mice were intravenously injected with live *C. albicans* (4×10^5^ cfu/100 μL 1×PBS), after *C. albicans* infected 5 days, mice kidney was isolated and followed by real-time PCR analysis of indicated proinflammatory genes expression (F) and acute kidney injury genes or adhesion molecules expression (G). Error bars represent SEM of three independent experiments for un-infected mice and six independent experiments for *C. albicans* infected mice. **(H)** Serum was isolated from the mice infected and treated as in (F). Urea and creatinine concentration were measured by ELISA analysis. Error bars represent SEM of three independent experiments for un-infected mice and six independent experiments for *C. albicans* infected mice. Two-tailed unpaired Student’s t test (C-H), two-way ANOVA (B), and log-rank (Mantel-Cox) test (A).

During fungal sepsis, host cell death could provoke inflammation and immunopathology and IL-23 signaling could prevent this process.^33^ Additionally, excessive inflammatory response can lead to serious collateral tissue damage and dysregulated immunopathology increases mice mortality during later stage of invasive candidiasis.^34^ So, we next explored whether the higher mortality of *Hilpda*^fl/fl^ Lyz2-Cre mice accompanied with higher inflammatory response and renal tissue damage. As speculated, we found the pro-inflammatory cytokines expression were significantly enhanced in *Hilpda*^fl/fl^ Lyz2-Cre mice’s kidney after *C. albicans* infection, including IL-6, IL-1β, TNF-α, IP-10, MCP1 and MIP1α (Figure 7F). These results are consistent with previous reports that high levels of IL-6, IL-1β, and TNF-α produced in kidney at initial stages of systemic *C. albicans* infection contributes to the immunopathogenesis of disease.^34–36^ Furthermore, the kidney injury molecule-1 (KIM-1) and neutrophil gelatinase-associated lipocalin (NGAL) were significantly up-regulated expression in *Hilpda*^fl/fl^ Lyz2-Cre mice after fungal infection (Figure 7G), a marker of tubular epithelial damage used for the early diagnosis of acute kidney injury.^37^ Meanwhile, the major adhesion molecules ICAM-1 and P-Selectin, which involved in mediating acute kidney injury,^38^ were also significantly increased in *Hilpda*^fl/fl^ Lyz2-Cre mice (Figure 7G). Consistent with the above results, the levels of serum urea and creatinine in *Hilpda*^fl/fl^ Lyz2-Cre mice were much higher than *Hilpda*^fl/fl^ mice during fungal infection (Figure 7H). Besides, we also explored the local neutrophils or macrophages recruitment in kidney from fungal infection mice. We found that there was no difference change of macrophage and neutrophil infiltration in the kidneys between *Hilpda*^fl/fl^ Lyz2-Cre and *Hilpda*^fl/fl^ mice after systemic *C. albicans* infection (Figure S8E and S8F). Th1 and Th17 cell differentiation play an important role in antifungal immunity as well. We further assessed whether *Hilpda* deficiency affected Th1 and Th17 responses after *C. albicans* infection. We found that Th1 and Th17 abundance was not markedly changed in the lymph nodes and spleen in *Hilpda*^fl/fl^ Lyz2-Cre mice compared with *Hilpda*^fl/fl^ mice after fungal infection (Figure S8G and S8H). Overall, these data indicate that LD plays an indispensable role in the antifungal immunity.

### ATGL inhibitor Atglistatin has a therapeutic impact on systemic fungal sepsis

Given our finding that LD accumulation is essential for phagolysosomes maturation and protects macrophages from death, we hypothesized that manipulating LD accumulation by ATGL inhibitors might augment antifungal immune responses *in vivo*. Utilizing the in *vivo* model of invasive candidiasis, we found that Atglistatin administration resulted *in* significantly prolonged survival compared to treatment with DMSO diluent alone (Figure 8A). We also observed decreased weight loss and reduced fungal burden in the kidneys of mice treated with Atglistatin (Figure 8B and 8C). Histopathological analysis and PAS staining also confirmed the reduction in fungal burden in the kidneys of mice treated with Atglistatin (Figure 8D).

**Figure 8.**
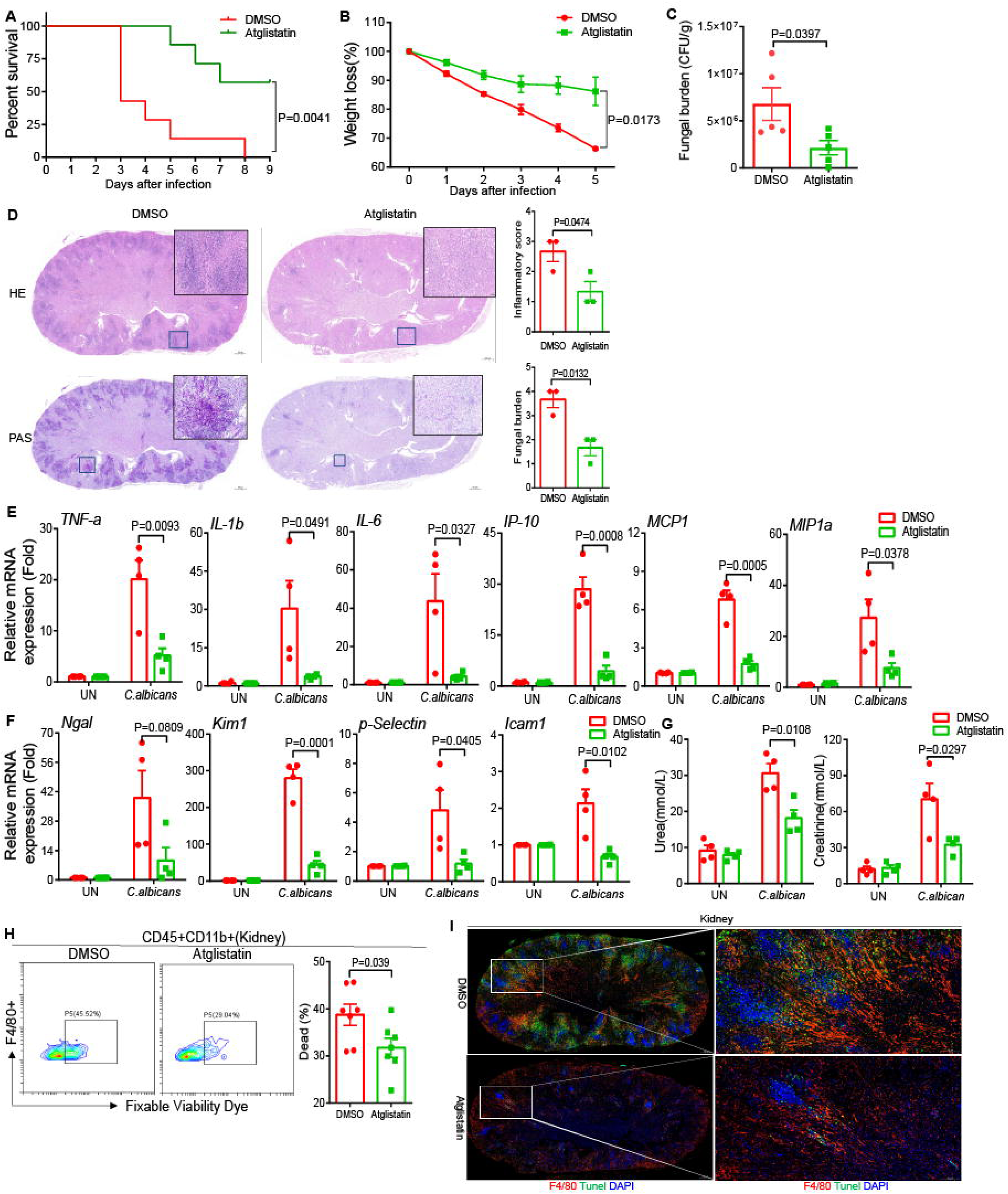
ATGL inhibitors treatment have a therapeutic impact on systemic fungal sepsis. **(A-C)** WT mice were intravenously injected with live *C. albicans* (1×10^5^ cfu/100μL 1×PBS) and intraperitoneally injected with DMSO or Atglistatin (10μg/50μL) after C. albicans infection. Survival rate (A), weight loss rate (B), or fungal number (C) in the kidney of mice were shown (n = 7 for A, n = 5 for B, and n = 5 for C). **(D)** WT mice were treated as in C, and 5 days after *C. albicans* infection, mice were euthanized, kidneys were fixed, and sections were stained with hematoxylin and eosin (H&E) or periodic-acid Schiff (PAS). Renal inflammation and fungal burden were scored based upon H&E and PAS staining (Right), respectively. Error bars represent SEM of three independent experiments. **(E-F)** WT mice were intravenously injected with live *C. albicans* (1×10^5^ cfu/100μL 1×PBS) and intraperitoneally injected with DMSO or Atglistatin (10μg/50μL), after *C. albicans* infected 5 days, mice kidney was isolated and followed by real-time PCR analysis of indicated proinflammatory genes expression (E) and acute kidney injury genes or adhesion molecules expression (F). Error bars represent SEM of four independent experiments. **(G)** Serum was isolated from the mice infected and treated as in (E). Urea and creatinine concentration were measured by ELISA analysis. Error bars represent SEM of four independent experiments. **(H)** WT mice were intravenously injected with live *C. albicans* (5×10^4^ cfu/100μL 1×PBS) and intraperitoneally injected with DMSO or Atglistatin (10μg/50μL), and mice were euthanized at day 5 after infection. The cell viability of macrophages from kidneys were stained for flow cytometry analysis. Error bars represent SEM of seven mice experiments. **(I)** WT mice were treated as in C, and 5 days after *C. albicans* infection, mice were euthanized, kidneys were fixed, and sections were stained with immunofluorescence. F4/80 indicated macrophages, TUNEL staining indicated dead cells and nuclei were stained with DAPI. Two-tailed unpaired Student’s t test (C-H), two-way ANOVA (B), and log-rank (Mantel-Cox) test (A).

We therefore explored whether Atglistatin executed its protective role of systemic fungal sepsis through inhibiting inflammatory response and tissue damage. We found the administration of Atglistatin after *C. albicans* infection decreased pro-inflammatory cytokines expression, including IL-6, IL-1β, TNF-α, IP-10, MCP1 and MIP1α (Figure 8E). Furthermore, Atglistatin treatment significantly down-regulated expression of kidney injury molecule-1 (KIM-1) and neutrophil gelatinase-associated lipocalin (NGAL) (Figure 8F). Meanwhile, the major adhesion molecules ICAM-1 and P-Selectin were also significantly reduced in Atglistatin treated mice (Figure 8F). Consistent with the above results, the levels of serum urea and creatinine in Atglistatin injected mice were much lower than control mice (Figure 8G). These further indicated that Atglistatin had a protective role on kidney from impairment after *C. albicans* infection. Besides, we also explored the local neutrophils or macrophages recruitment in kidney from fungal infection mice. We found that there was no difference change of macrophage and neutrophil infiltration in the kidneys between Atglistatin and control DMSO treated mice after systemic *C. albicans* infection (Figure S9A and S9B). However, the cell death of macrophages infiltration in the kidneys was significantly declined after treatment with Atglistatin (Figure 8H), and TUNEL staining also showed that the cell death of renal tissue cells was reduced by Atglistatin administration, including infiltrated macrophages (Figure 8I). We further assessed whether Atglistatin treatment regulated Th1 and Th17 responses after *C. albicans* infection. We found that Th1 and Th17 abundance was not markedly changed in the lymph nodes and spleen in Atglistatin treatment mice compared with control mice after fungal infection (Figure S9C and S9D). Meanwhile, we found that administration of Atglistatin could increase LD production in kidneys after *C. albicans* infection (Figure S9E). Overall, these data indicate that the ATGL is a therapeutic target for the treatment of systemic fungal sepsis, and inhibitors of ATGL have the potential to be developed as drugs for the treatment of invasive fungal infection.

## DISCUSSION

Cell metabolism is crucial for activating effective immune responses to resist invading pathogens. In fungal-infected macrophages, the switch from oxidative phosphorylation to glycolysis helps enhance immune function. Supplementing glucose during such infections can protect macrophages and aid mice.^18^ Neutrophils increase glucose intake by boosting Glut1 expression via activated PKC-δ, supporting their fungicidal activity.^39^ Additionally, 2-deoxy-D-glucose reduces zinc levels in cells, targeting the fungal pathogen *Histoplasma capsulatum.*^40^ Research is limited on other metabolic pathways, like lipid metabolism. Our research filled the knowledge gap and revealed that lipid metabolism seems involved in antifungal responses as fungal infections cause an increase in macrophage triglyceride levels and lipid droplets accumulation, likely due to increased lipid synthesis and reduced degradation.

Our research further demonstrated that increased triglycerides and LD accumulation in macrophages can limit phagosome formation and prevent cell death from excessive phagocytosis, which fails to clear fungi effectively, thus protecting the cells. This led to a hypothesis that macrophages should engulf fungi optimally within their capacity for effective elimination. Excessive phagocytosis, harmful in other conditions like atherogenesis where it can worsen disease progression by promoting foam cell formation, shows similar patterns in different contexts.^41^ Additionally, reports show that ATGL deletion in macrophages leads to LD accumulation and decreased phagocytic ability, attributed to reduced ATP levels.^42^ Conversely, our findings suggest that LD accumulation during fungal infection hampers phagocytosis by occupying the ER components and sequestering RAC1 on LD surfaces, thus reducing excessive phagocytosis (Figure S10).

Traditional models of phagocytosis describe the plasma membrane as the primary source for phagosome formation through invagination.^43,44^ However, contributions from other endo-membranes, notably the ER, in what’s termed ER-mediated phagocytosis, are also significant.^12^ This involvement is evidenced by the presence of both plasma membrane and ER proteins (like calreticulin and calnexin) on phagosomes.^45,46^ Phagosomes typically undergo sequential fusion with early endosomes, late endosomes, and lysosomes.^15,16^ We previously identified that ER-derived membranes can supplement phagosome membranes, with the ER protein STING translocation to phagosomes during fungal stimulation.^17^ Additionally, ER membranes marked by calnexin transfer to phagosomes has also been observed in this study, suggesting complex interactions between the ER and phagosome formation that require further investigation.

During disseminated fungal infection, the high fungal load in kidney and late renal immunopathology were the two main reasons leading to mortality. The high levels of IL-6, IL-1β, and TNF-α produced in kidney of systemic *C. albicans* infection contributes to the immunopathogenesis of disease.^34–36^ Our data showed that administration of ATGL inhibitors Atglistatin not only diminished the fungal burden, but also could effectively alleviate inflammatory reaction and reduce kidney injury. The diminished fungal burden was due to Atglistatin induced LD accumulation protected macrophages from death and enhanced macrophages’ fungicidal ability. In addition, intracellular accumulated LD could suppress inflammatory reaction in immune cell. For example, LPS-mediated activation markedly increased LD accumulation or the reduction in ATGL-mediated lipolysis attenuated the inflammatory response in macrophages, which was accompanied by decreased production of prostaglandin-E2 (PGE2) and IL-6.^47^

Currently, two primary strategies exist for developing antifungal therapies. Firstly, small molecule drugs like Azole, and Echinocandin target fungal enzymes to inhibit vital components of fungal growth, ^48,49^ but these drugs often cause severe side effects and lead to drug resistance. To address resistance, researchers discovered Turbinmicin from *Micromonospora sp*, which has shown effectiveness against multi-drug-resistant fungal pathogens like *Candida auris.* ^50^ Secondly, there is a focus on enhancing host antifungal immune responses through small molecules or other therapies. Recent studies have explored the potential of JNK1 inhibitors, Sirt2 deacetylase inhibitors (AGK2, AK-1, AK-7), ROS activators, and nanoparticle-mediated delivery of Rac1 mRNA to boost antifungal immunity.^51, 52, 31^ Furthermore, we found the negative role of STING in antifungal immunity, suggesting its N-terminal peptide as a potential therapeutic target.^17^ In this study, we found ATGL inhibitors like Atglistatin is effective in protecting mice from systemic *C. albicans* infections, indicating ATGL as a promising drug target. Future plans include synthesizing small molecule peptides based on the HILPDA-ATGL interaction to assess their therapeutic effects in systemic fungal sepsis models, aiming to develop safer and more effective antifungal treatments.

## ACKNOWLEDGMENTS

We thank Dr. Changbin Chen (Institute Pasteur of Shanghai, CAS, Shanghai, China) for kindly providing *C. albicans* strain SC5314. This work was supported by the grants from the National Natural Science Foundation of China (82321002, 32230033 and 81930039 to C.G.), National key research and development program of China (2021YFC2300603 and 2023YFC2306102 to C.G.). This work also supported by the Future Scholar Program of Shandong University (21510082364125).

## AUTHOR CONTRIBUTIONS

C.G. and W.S. conceived and designed the study. W.S. and H.W. performed the experiments with the assistance from G.Z., Q.S., L.Z., X.L., and T.C., F.L., Y.Z., W.Z., X.Q., B.L. contributed to the discussion and provided reagents. C.G. and W.S. analyzed the data and wrote the paper, and T.C. helped to polish the English writing.

## Materials and methods

### Mice

The *Hilpda*^fl/fl^ mice were constructed by GemPharmatech using the CRISPR-Cas9gene-editing system. Lyz2-Cre mice were obtained from Jackson Laboratory. *Hilpda*^fl/fl^ mice were crossed with Lyz2-Cre mice to generate *Hilpda*^fl/fl^ Lyz2-Cre mice. The Genotyping of *Hilpda*^fl/fl^ was confirmed by means of PCR using the following primers: forward primer 5’-TCTGAGGCGGAAAGAACCAG-3’ and reverse primer 5’-TGAACTCCTCCATCTTTCAGCC-3 for identifying 5’arm, forward primer 5’-TAGGGCTGCATAGCAAGACTGAAC-3’ and reverse primer 5’-CTGCAAAGGAC-CCAACTGTATG-3 for identifying 3’arm. The primers for Lyz2-Cre transgenic mice genotyping were oIMR3066 mutant 5’-CCCAGAAATGCCAGATTACG-3’, oIMR3067 common 5’-CTTGGGCTGCCAGAATTTCTC-3’ and oIMR3068 WT 5’-TTACAGTCGGCCAGGCTGAC-3’. To avoid possible off-target editing of the mice genome, the mice were bred six generation with mice of C57BL/6 background before performing any experiments. Mice were housed under SPF condition, and the experimental protocols were based on the general guideline principles provided by the Association for Assessment and Accreditation of Laboratory Animal Care, and performed with approval from Scientific Investigation Board of the Medical School of Shandong University.

### Cells

293T and Hela cells were maintained in DMEM plus 10% FBS and 1% Penicillin-Streptomycin. THP1 and RAW264.7 cells were maintained in RPMI-1640 plus 10% FBS and 1% Penicillin-Streptomycin and 1% sodium pyruvate.

### Mouse BMDMs preparation

BMDMs were obtained by differentiating bone marrow progenitors from the tibia and femur of 6–8-week-old male or female mice in Iscove’s Modified Dulbecco’s Media (IMDM) containing 20ng/mL of M-CSF, 10% heat-inactivated fetal bovine serum (FBS, Invitrogen), 1mM sodium pyruvate, 100 U/mL penicillin, and 100μg/mL streptomycin (Invitrogen) for 5-7 days. Cells were then re-plated in 6-well or 12-well plates 1 day before experiments.

### Reagents

Antibodies of anti-ATGL (clone 30A4), anti-p-Akt (Ser473) (clone D9E), anti-p-Akt (Thr308) (clone D25E6), and anti-Rab7 (clone D95F2) were bought from Cell Signaling Technology (cat no. 2439, 4060, 13038 and 9367). Antibodies of anti-HA and anti-Flag for immunoprecipitation were bought from Origene (cat no. TA100012) and Sigma-Aldrich (cat no. F1804), respectively. Antibodies of anti-PLIN2 (clone G-2), anti-PLIN3 (clone B-3), anti-HILPDA (clone G-2), anti-GAPDH (clone 2E3-2E10), anti-Tom20 (clone F-10) and anti-HSP90 (clone F-8) were bought from SANTA CRUZ BIOTECHNOLOGY (cat no. sc-390169, sc-390981, sc-376704, sc-293335, sc-17764 and sc-13119). Antibody of anti-Calnexin, anti-GM130, and anti-RAC1 was bought from Proteintech (cat no. 10427-2-AP, 11308-1-AP and 66122-1-Ig). Antibody of anti-Syk was bought from Enzo Life Sciences (cat no: ALX-804-480-C100). Antibody of anti-FcεRI antibody, γ subunit was bought from Millipore (cat no: 06-727). Flow antibodies of anti-CD45 (clone 30-F11), anti-Ly6G (clone 1A8), anti-F4/80 (clone BM8), anti-CD4(clone GK1.5), anti-IL-17A (clone TC11-18H10.1) and anti-CD11b (clone M1/70) were bought from Biolegend (cat no: 103108, 127608, 123115, 100406, 506908 and 101212). Antibody of anti-IFN-γ (clone XMG1.2) was bought from eBioscience (cat no.11-7311-82). Zombie Violet Fixable Viability kit was bought from Biolegend (cat no. 423114). ZymosanA-Alexa Fluor™ 594, D-zymosan and curdlan were bought from Invivogen (cat no. Z23374, tlrl-zyd and tlrl-cud). A922500, T863 and Atglistatin were bought from MCE (HY-10038, HY-32219 and HY-15859). Recombinant mouse M-CSF proteins were bought from Peprotech (cat no. 315-02).

### Immunofluorescence

Cells were fixed with 4% paraformaldehyde and followed by permeabilization treatment with 1×PBS containing 0.3% Triton X-100 for 10 minutes. Prior to incubation with primary antibody, samples were incubated with 10% goat serum at room temperature for 1h to block non-specific staining. After 12h of incubation with primary antibody at 4°C, the samples were washed three times with ice cold 1×PBS and further stained with fluorophore (Alex 488 or Alex 405 and Alexa 568 conjugated secondary antibodies). After staining, samples were counter stained with DAPI and immersed in mounting medium before being sealed on a slide with nail polish. Sealed slides were analyzed using high-speed confocal microscopy (Andor Dragonfly 200) with companion software.

### Immunoblot and immunoprecipitation

Cells were harvested and lysed on ice in lysis buffer which containing 0.5% Triton X-100, 20 mM Hepes pH 7.4, 150 mM NaCl, 12.5 mM β-glycerophosphate, 1.5mM MgCl2, 10mM NaF, 2mM dithiothreitol, 1mM sodium orthovanadate, 2mM EGTA, 20 mM aprotinin, and 1mM phenylmethylsulfonyl fluoride for 30 minutes, followed by centrifuging at 12,000 rpm for 15 minutes to extract clear lysates. For immunoprecipitation, cell lysates were incubated with 1μg of antibody at 4°C overnight, followed by incubation with A-sepharose or G-sepharose beads for 2h, and the beads were washed four times with lysis buffer and the precipitates were eluted with 2×sample buffer. Elutes and whole cell extracts were resolved on SDS-PAGE followed by immunoblotting with antibodies.

### Lentivirus-medicated gene knockout/knockdown in THP1 or RAW264.7 cells

pLentiCRISPR-GFP vector was used for CRISPR/Cas9-mediated gene knockout in THP1 or RAW264.7 cell line. Briefly, lentivirus vector expressing gRNA was transfected together with package vectors into HEK293T (ATCC) package cells. 48 and 72 h after transfection, virus supernatants were harvested and filtrated with 0.2μm filter. Target cells were infected twice and sorted by flow cytometry medicated cell sorting. For some experiments, single cell was plated into 96-well plate by flow cytometry for single clone isolation. Isolated single clones were verified by western blot and DNA sequencing.

### RT and Real-time PCR

Total RNA was extracted from spinal cord with TRIzol according to the manufacturer’s instructions. 1μg total RNA for each sample was reverse transcribed using the SuperScript® II Reverse Transcriptase from Thermo Fisher Scientific. The resulting complementary DNA was analyzed by real-time PCR using SYBR Green Real-Time PCR Master Mix. All gene expression results were expressed as arbitrary units relative to expression β-*actin* or *GAPDH*.

### Flow cytometry

For *in vitro* experiments, THP1, RAW264.7 or BMDMs cells were stimulated with HKCA, D-zymosan or Curdlan followed by staining with BODIPY or Nile-Red for marking LD. Then cells were analyzed by flow cytometry. For *in vivo* experiments, 5 days after live *C. albicans* infection, mice were sacrificed and perfused with 1×PBS. Kidneys were homogenized in ice cold tissue grinders, filtered through a 100μm cell strainer and the cells collected by centrifugation at 400g for 5 minutes at 4°C. Cells were resuspended in 10mL of 30% Percoll (Amersham Bioscience) and centrifuge onto a 70% Percoll cushion in 15-mL tubes at 800g for 30 minutes. Cells at the 30-70% interface were collected and were subjected to flow cytometry. Cell surface staining was done for 30 minutes at 4°C. Zombie Violet Fixable Viability kit (1:400; Biolegend) was added to exclude dead cells. Flow cytometry data analysis was performed by using CytExpert.

### BMDMs phagocytosis assay

For BMDMs phagocytosis assays, *C. albicans* (strain SC5314) cells were labeled with Alexa Fluor 488 in 100 mM HEPES buffer (pH 7.5). BMDMs were co-cultured with labeled *C. albicans* at 37 °C for 45 minutes. Fluorescence signals from fungal cells that were adherent to, but not phagocytosed by, phagocytes were quenched with trypan blue, and the rate of phagocytosis was assessed by flow cytometry. For GFP-*C. albicans*, funguses were fixed by 2% paraformaldehyde at room temperature for 30 minutes, then were co-cultured with BMDMs for the indicated times. Unbound yeasts were gently washed by 1×PBS with five times, cells were analyzed by flow cytometry.

### Cell subpopulation of BMDMs phagocytosis

HKCA was dual labeled by FITC and Fluor-647, and then analyzed the cell subpopulation which contained phagosomes of different numbers according to “Dendritic Cell Protocols” edited by Shalin H. Naik.^30^ Briefly, a total of 1ⅹ10^6^ BMDMs or RAW264.7 cells were seeded in 6-cm plates. Cells were infected with adequate FITC and Fluor-647 labeled HKCA (MOI=10) for the 1h, 2h or 4h, then added 2mL of ice-cold 1×PBS to stop the internalization and remove the non-internalized HKCA for 5 times. Added 1mL conditioned-complete medium and incubated at 37 °C and 5% CO_2_ for another 2h. Cells were harvested and analyzed by flow cytometry.

### RAC1 activity assay

A total of 5×10^6^ THP1 cells or RAW264.7 and DGAT1-KO RAW264.7 cells were seeded in 6-cm plates. After THP1 cells were pretreated with OA (4.5ug/mL), A922500 (10uM), or T863 (10uM) for 24h, cells were infected for 0, 30 and 60 minutes with HKCA (MOI=3). Cells were washed twice with ice-cold 1×PBS and lysed in lysis buffer containing 50mM Tris (pH 7.5), 10mM MgCl_2_, 0.5M NaCl, and 2% IGEPAL, as described in the Rac1/Cdc42 Activation Assay Kit manual (Upstate). The supernatants were incubated with GST-PBD-agarose beads at 4°C for 4h, washed three times in wash buffer (25mM Tris (pH 7.5), 30mM MgCl_2_, 40mM NaCl) and resuspended in loading buffer for immunoblot analysis.

### Thin layer chromatography assay

Cells were scraped from a 10-cm plates, then added 2mL 1×PBS containing protease inhibitor and 1mM PMSF and 4mL chloroform/methanol (2:1, vol/vol) and 0.01% butyl hydroxytoluene to obtain the total lipid extraction. Vortex the solution at a rate of 1500×g for 5 minutes. Collect organic phase, add 2.5mL of chloroform to the aqueous phase to extract remaining lipids, vortex, 1500×g for 5 minutes. Drop the total lipid extraction solution dissolved in chloroform onto a 5-10 cm EMD thin-layer chromatography silica gel plate, soak the bottom with hexane/ether/acetic acid (80:20:2, vol/vol/vol) as the spreading agent, and allow the thin-layer chromatography silica gel plate to absorb the spreading agent before developing. Observe the lipids using iodine vapor.

### Cell full spectrum metabolome

For the metabolomic experiment, BMDMs were isolated from wild type mice. Cells were treated with D-zymosan (20μg/mL) for 4h and were harvested for metabolites extraction. Samples were sent to the Metware company for metabolomic analysis. Each time point had six parallel replications. For extraction of hydrophilic compounds, sample was thawed on ice, then added 1mL pre-cooled extractant (70% methanol aqueous solution), and whirl for 1 minute. Freeze the mixture for 3 minutes in liquid nitrogen after remove ice for 3 minutes, it will be whirled for 2 minutes, circulate this at 3 times. Centrifuge the mixture again with 12,000 rpm/min at 4℃ for 10 minutes. Finally take the supernatant into the sample bottle for LC-MS/MS analysis. For extraction of hydrophobic compounds, sample placed in liquid nitrogen for 2 minutes then thawed on ice for 5 minutes and vortex blending. Repeat the first step 3 times, then centrifuge it with 12,000 rpm at 4℃ for 10 minutes. Take 300uL supernatant and homogenize it with 1mL mixture (include methanol, MTBE and internal standard mixture). Whirl the mixture for 2 minutes. Then add 500μL of water and whirl the mixture for 1 minute, and centrifuge it with 12,000 rpm at 4℃ for 10 minutes. Extract 500μL supernatant and concentrate it. Dissolve powder with 100μL mobile phase B, then stored in −80℃. Finally take the dissolving solution into the sample bottle for LC-MS/MS analysis.

### RNA-seq

For the RNA-seq experiment, BMDMs were isolated from wild type mice. Cells were treated with Curdlan (20μg/mL) for 0, 3 and 6h, and were harvested by Trizol. Samples were sent to the BGI company for RNA-seq analysis. Each time point had three parallel replications. For the data analysis, heatmap of changes in gene expression of selected genes in stimulated macrophages over non-stimulated

### Mass spectrometry identification of LD proteins

For the identification of LD surface proteins, LD was isolated from THP1 cells which pretreated with OA and then stimulated with or without HKCA for 4h. The specific experimental process was according to instruction of Lipid Droplet Extraction Kit (ab242290). After LD was lysed, the released proteins were analyzed by mass spectrometry analysis.

Samples were reduced and alkylated in dithiothreitol (DTT) and iodoacetamide followed by trypsin digestion overnight. Digested samples were injected onto Agilent Zorbax 300SB-C18 0.075mm×150mm column on Eskigent nano LC system coupled with Thermo LTQ-ETD-Orbitrap. Advion Triversa nanomate served as the nano-ion spray source. MSMS data were searched against Refseq human protein database by Sorcerer Sequest. The searched dataset was processed by TPP (Trans-Proteomics Pipeline) and filtered with Peptide Prophet.

### Lactate dehydrogenase cytotoxicity assay

LDH release was assayed using the Pierce™ LDH Cytotoxicity Assay Kit (Thermo Fisher Scientific) following the manufacturer’s protocol. Culture medium was collected and centrifuged at 400×g for 5 minutes to remove cell debris. LDH release into the medium was measured at OD490. Relative LDH release was expressed as the percentage LDH activity in supernatants of cultured cells (medium) compared to total LDH (from media and cells) and used as an index of cytotoxicity.

### Systemic *C. albicans* infection model

Live *C. albicans* strain SC5314 (4×10^5^ yeast cells in 0.1mL of 1×PBS buffer) were injected intravenously into 6- to 8-week-old littermates of distinct genotypes. Infected mice were monitored daily for weight loss and survival. Fungal burden of kidneys was measured 5 days after infection. After the kidneys were collected, tissue homogenates were serially diluted and plated on yeast extract–peptone–dextrose agar. Fungal colony-forming units were counted 24h after plating. For the mice treatment of Atglistatin experiments, Atglistatin (20μg/mouse) was intraperitoneal injected 12h after *C. albicans* infection (intravenously injection of 1×10^5^ live *C. albicans* strain SC 5314 yeast cells in 0.1mL of 1×PBS), and weight loss and survival rate were calculated.

### Histopathology

For histopathology analyses, kidneys were fixed in 10% neutral-buffered formalin, processed according to standard procedures, embedded in paraffin, and sectioned. 2μm-thick sections were stained with hematoxylin and eosin (H&E), periodic-acid-Schiff (PAS). Renal inflammation was scored based upon H&E and PAS staining (proportion of renal parenchyma and/or pelvis involved by tubulointerstitial nephritis and/or pyelonephritis) as not significant (score 0), less than 10% (score 1), 10-25% (score 2), 25-50% (score 3), or greater than 50% (score 4). The intra-lesional fungal burden was based upon PAS staining as not significant (score 0), scant presence in less than 10% of inflammatory foci (score 1), mild-to-moderate presence in 10-25% of inflammatory foci (score 2), moderate-to-significant presence in 25-50% of inflammatory foci (score 3), or significant presence in more than 50% of inflammatory foci (score 4).

### Statistics

Non-parametric statistics was applied to compare differences between two groups. Two-tailed unpaired t test was used to derive all of the *P* value; the clinical scores and weight change curve were analyzed by two-way ANOVA for multiple comparisons. *P <*0.05 was considered to be significant. Results are shown as mean and the error bar represents standard error of mean (S.E.M) biological replicates as indicated in the figure legend.

## Supplemental Information

### Supplementary Figure legend

**Figure S1**. **Metabolomics profiling analyzed the changed metabolites in BMDMs stimulated with D-zymosan.**

**(A)** Fold change (log2) of all differential metabolites in D-zymosan-stimulated mice BMDMs over non-stimulated macrophages.

**(B)** Principal component analysis showing distinct clustering of different treatments.

**(C)** Volcano plot describing the abundance of differentially metabolites. Two vertical dashed lines on each side correspond to −1.0 and 1.0 cut-points, which are Log2(fold change) cutoffs used for determining differentially metabolites. The horizontal dashed line corresponds to P-adjusted values of 0.05, which was another cutoff used for determining differentially metabolites. Arrow indicated up-regulated TGs.

**(D)** Enrichment analysis of signaling pathways for differential metabolites. Arrow indicated lipid metabolism related signaling pathways.

**(E)** TLC analysis and quantification of triglycerides (TG) levels in BMDMs stimulated with different fungal ligands. Error bars represent SEM of three biological replicates.

**(F)** TLC analysis and quantification of triglycerides (TG) levels in RAW264.7 cells stimulated with D-zymosan (100μg/mL) for 4h. Error bars represent SEM of three biological replicates.

**(G)** TLC analysis and quantification of triglycerides (TG) levels in THP1 cells stimulated with different fungal ligands. Error bars represent SEM of three biological replicates.

**(G)** Quantification of triglycerides (TG) levels in BMDMs stimulated with HKCA (MOI=3) for the indicated times.

**(H)** Quantification of triglycerides (TG) levels in THP1 cells stimulated with HKCA (MOI=3) for the indicated times.

Two-tailed unpaired Student’s t test (E, F and G) and one-way ANOVA for (H and I).

**Figure S2. The RNA-seq data of BMDMs stimulated with curdlan for different times was validated.**

**(A)** Heatmap of changes in gene expression of lipid biogenesis and lipolysis genes in *C. albicans*-stimulated macrophages over non-stimulated macrophages from Timothy et al. (2018).

**(B)** BMDMs were stimulated with HKCA (MOI=3), D-zymosan (100μg/mL) and curdlan (100μg/mL) for the indicated times followed by real-time PCR analysis of genes in indicated functional classes. Error bars represent SEM of three biological replicates.

**(C)** BMDMs were stimulated with HKCA (MOI=3), D-zymosan (100μg/mL) and curdlan (100μg/mL) for the indicated time followed by Western blot analysis of LD structural proteins PLIN2 and PLIN3. Representative immunoblot images are shown.

**(D)** DGAT1 or HILPDA knockout cell lines were generated using the CRISPR-Cas9 system, verified by Western blot.

**(E)** THP1 cells were pretreated with A922500 (10μM) or T863(10μM) for 24h followed by stimulation with HKCA (MOI=3) for 4h. LD were analyzed by immunofluorescence staining with BODIPY and nucleus was stained by DAPI (Scale bar: 40μm), and LD-BODIPY MFI was measured by flow cytometry (Right). Error bars represent SEM of three biological replicates.

**(F)** THP1 cells were pretreated with Atglistatin (10μM) for 24h followed by stimulation with HKCA (MOI=3) for 4h. LD were analyzed by immunofluorescence staining with BODIPY and nucleus was stained by DAPI (Scale bar: 40μm), and LD-BODIPY MFI was measured by flow cytometry (Right). Error bars represent SEM of three biological replicates.

Two-tailed unpaired Student’s t test (E and F) and one-way ANOVA for (B).

**Figure S3. LD formation had no effect on phagocytosis.**

**(A)** Phagocytosis of THP1 cells treated with or without A922500 (10μM), T863 (10μM) and Atglistatin (10μM), control THP1 cells or DGAT1-KO THP1 cells, control RAW264.7 cells or DGAT1-KO RAW264.7 cells, control THP1 cells or HILPDA-KO THP1 cells, and *Hilpda*^fl/fl^ or *Hilpda*^fl/fl^ Lyz2-Cre mice BMDMs was evaluated by the method described in Materials and Methods. Error bars represent SEM of three or four biological replicates.

**(B-C)** BMDMs(B) or THP1 cells(C) were pretreated with OA (4.5μg/mL) for 24h, followed by immunofluorescence staining. LD was stained by BODIPY and nucleus was stained by DAPI. (Scale bar: 20μm for B and 40μm for C). Meanwhile, LD-BODIPY MFI was measured by flow cytometry. Error bars represent SEM of three biological replicates.

**(D)** THP1 cells were pretreated with different concentration of OA (4.5μg/mL) for 24h, followed by flow cytometry to test LD-BODIPY.

**(E)** Phagocytosis of THP1 cells were pretreated with different concentration of OA for 24h was evaluated by the method described in Materials and Methods. Error bars represent SEM of three biological replicates.

Two-tailed unpaired Student’s t test (A, B and C) and one-way ANOVA for (D and E).

**Figure S4. LD accumulation attenuated phagosomes formation and numbers.**

**(A-C)** THP1 cells were pretreated with or without OA (4.5μg/mL, A), Atglistatin (10μM, B), A922500 and T863(10μM, C) for 24h and then engulfed HKCA-GFP (MOI=10) for the indicated times followed by immunofluorescence staining of indicated proteins (Scale bar: 40μm and 10μm for representative single cell images). Nuclei were stained with DAPI and Syk staining indicated cell position. The number of phagosomes containing HKCA-GFP was counted in single cell (lower). N represented cell number.

**(D-F)** BMDMs cells were pretreated with or without OA (4.5μg/mL, D), Atglistatin (10μM, E), A922500 and T863 (10μM, F) for 24h and then engulfed FITC and Fluor-647 double labeled HKCA (MOI=10) for 30 minutes and digested for another 2h followed by flow cytometry. A detailed analysis of the phagocytic population in the same plot makes evident the presence of different cell populations according to the number of phagosomes internalized by an individual cell (population a: cells with a small number of phagosomes; population b: cells with quite a few phagosomes; population c: cells with moderate number of phagosomes; population d: cells with a large number of phagosomes). Error bars represent SEM of six biological replicates.

Two-tailed unpaired Student’s t test (A-F).

**Figure S5. Phagosome formation affected LDs accumulation.**

**(A-B)** Mouse macrophages were stimulated with different infection ratio of HKCA followed by immunofluorescence staining of indicated proteins (A) (Scale bar: 10μm) and flow cytometry to detect the LD-BODIPY MFI (B). Calnexin indicated endoplasmic reticulum, CFW stained fungal cell wall and LD was stained by BODIPY. Error bars represent SEM of three biological replicates.

**(C)** Phagocytosis of THP1 cells treated with Ly294002 (10μM), SL-2052 (10μM) and XL-147 (10μM) was evaluated by the method described in Materials and Methods. Error bars represent SEM of three biological replicates.

**(D)** BMDM cells were pretreated with SL-2052 (10μM) for 1h, then stimulated with HKCA (MOI=3) for the indicated time followed by Western blot analysis of indicated proteins expression. Representative immunoblot images are shown.

**(E)** BMDM cells were pretreated with SL-2052 (10μM) for 1h, then stimulated with HKCA (MOI=3) for 4h followed by real-time PCR analysis of indicated gene expression. Error bars represent SEM of three biological replicates.

Two-tailed unpaired Student’s t test (C and E) and one-way ANOVA for (B).

**Figure S6. LD accumulation had no effect on PI3K/Akt activation and LD surface proteins.**

**(A-C)** THP1(A), RAW264.7(B) and BMDM (C) cells were pretreated with OA (4.5μg/mL) for 24h, then stimulated with HKCA (MOI=3) for the indicated time followed by Western blot analysis of indicated proteins. Representative immunoblot images are shown.

**(D)** The colocalization of LD and RAC1 was analyzed by immunofluorescence (Scale bar: 10μm). LD was stained by BODIPY.

**(E)** The colocalization of lipid droplet structural protein PLIN1 and RAC1 was analyzed by immunofluorescence (Scale bar: 10μm).

**(F)** THP1 cells were pretreated with OA (4.5μg/mL) for 24h, then intracellular lipid droplets were isolated and Western blot analysis of indicated proteins. Representative immunoblot images are shown.

**(G)** THP1 cells were stimulated with HKCA (MOI=3) for the indicated time, then intracellular lipid droplets were isolated and Western blot analysis of indicated proteins. Representative immunoblot images are shown.

**Figure S7. LD accumulation reduced the death of macrophages after *C. albicans* infection.**

**(A)** THP1 cells were pretreated with or without OA (4.5μg/mL) for 24h, then were infected with live hyphae-deficient (cph1Δ/Δ, efg1Δ/Δ) *C. albicans* (MOI=10) for the indicated time followed by PI staining, and dying THP1 cells were stained with PI (red) (Scale bar: 20μm).

**(B-C)** THP1 cells were pretreated with or without Atglistatin (10μM) for 24h, then were infected with live wild-type *C. albicans* or hyphae-deficient (cph1Δ/Δ, efg1Δ/Δ) C. albicans (MOI=10) for the indicated time followed by a lactate dehydrogenase (LDH) cytotoxicity assay of THP1 cells (B), and dying THP1 cells were stained with PI (red) (Scale bar: 20μm, C). Error bars represent SEM of three biological replicates.

**(D)** BMDMs cells were pretreated with or without Atglistatin (10μM) for 24h, then were infected with live wild-type *C. albicans* for 9h followed by flow cytometry to test cell viability. Error bars represent SEM of five biological replicates.

**(E-F)** THP1 cells were pretreated with DMSO, A922500 (10μM) or T863 (10μM) for 24h, then were infected with live wild-type *C. albicans* or hyphae-deficient (cph1Δ/Δ, efg1Δ/Δ) *C. albicans* (MOI=10) for the indicated time followed by a lactate dehydrogenase (LDH) cytotoxicity assay of THP1 cells (E), and dying THP1 cells were stained with PI (red) (Scale bar: 20μm, F). Error bars represent SEM of three biological replicates.

**(G)** BMDMs cells were pretreated with DMSO, A922500 (10μM) or T863 (10μM) for 24h, then were infected with live wild-type *C. albicans* for 9h followed by flow cytometry to test cell viability. Error bars represent SEM of four biological replicates. Two-tailed unpaired Student’s t test (B, D, E and G) and one-way ANOVA for (C and F).

**Figure S8. HILPDA interacts with ATGL.**

**(A-B)** HEK293T cells were transfected with indicated plasmids, and cell lysates were immunoprecipitated with anti-Flag antibody (A) or anti-HA (B) antibody followed by immunoblot analysis for indicated proteins. Asterisk indicated IgG light chain. Representative immunoblot images are shown.

**(C)** Hela cells were transfected with ATGL-Flag together with HILPDA-HA WT or different deletion mutants, followed by immunofluorescence staining of anti-Flag and anti-HA antibodies (Scale bar: 10μm).

**(D)** *Hilpda*^fl/fl^ or *Hilpda*^fl/fl^ Lyz2-Cre BMDM cells were stimulated with HKCA (MOI =3) for the indicated time followed by Western blot to examine HILPDA knockout efficiency.

**(E-F)** *Hilpda*^fl/fl^ or *Hilpda*^fl/fl^ Lyz2-Cre mice were intravenously injected with live *C. albicans* (4×10^5^ cfu/100μL 1×PBS), and mice were euthanized at day 5 after infection. Neutrophiles(E) and macrophages(F) from kidneys were stained for flow cytometry analysis. Error bars represent SEM of seven mice experiments.

**(G-H)** *Hilpda*^fl/fl^ or *Hilpda*^fl/fl^ Lyz2-Cre mice were intravenously injected with live *C. albicans* (5×10^4^ cfu/100μL 1×PBS), and mice were euthanized at day 7 after infection. Cells from lymph nodes (G) and spleen (H) were stained for flow cytometry analysis. Error bars represent SEM of seven mice experiments.

Two-tailed unpaired Student’s t test (E-H).

**Figure S9. Immune cell infiltration and T helper subsets differentiation after administration with Atglistatin during fungal sepsis.**

**(A-B)** WT mice were intravenously injected with live *C. albicans* (5×10^4^ cfu/100μL 1×PBS) and intraperitoneally injected with DMSO or Atglistatin (10μg/50μL), and mice were euthanized at day 5 after infection. Neutrophiles(A) and macrophages(B) from kidneys were stained for flow cytometry analysis. Error bars represent SEM of seven mice experiments.

**(C-D)** WT mice were intravenously injected with live *C. albicans* (5×10^4^ cfu/100μL 1×PBS) and intraperitoneally injected with DMSO or Atglistatin (10μg/50μL), and mice were euthanized at day 7 after infection. Cells from lymph nodes (C) and spleen (D) were stained for flow cytometry analysis. Error bars represent SEM of seven mice experiments.

**(E)** WT mice were treated as in C, and 5 days after *C. albicans* infection, mice were euthanized, kidneys were fixed, and sections were stained with immunofluorescence. F4/80 indicated macrophages, LD was stained by BODIPY and nuclei were stained with DAPI.

Two-tailed unpaired Student’s t test (A-D).

**Figure S10. LDs control phagosomes formation excessively and promote phagolysosomes formation during *C. albicans* infection.**

Upon *C. albicans* infection, intracellular LDs were accumulated gradually and the accumulation was induced by lipid biosynthesis increasement and inhibited lipolysis. On one hand, LD accumulation competitively consumed intracellular endoplasmic reticulum membrane components, which were also exploited by phagosomes formation. On the other hand, massive LD formation recruited Rho family small GTP enzymes, RAC1/2, to LDs’ surface. This alteration of RAC1 location resulted in inactivation of RAC1 GTPases activity, which was important for phagocytosis through promoting microfilaments aggregation. In addition, LD formation also recruited endoplasmic transport protein-Rab7 and then enhanced the fusion between phagosomes and lysosomes. These processes promoted phagolysosomes formation and improved macrophages survival.

**Supplementary Table S9:**
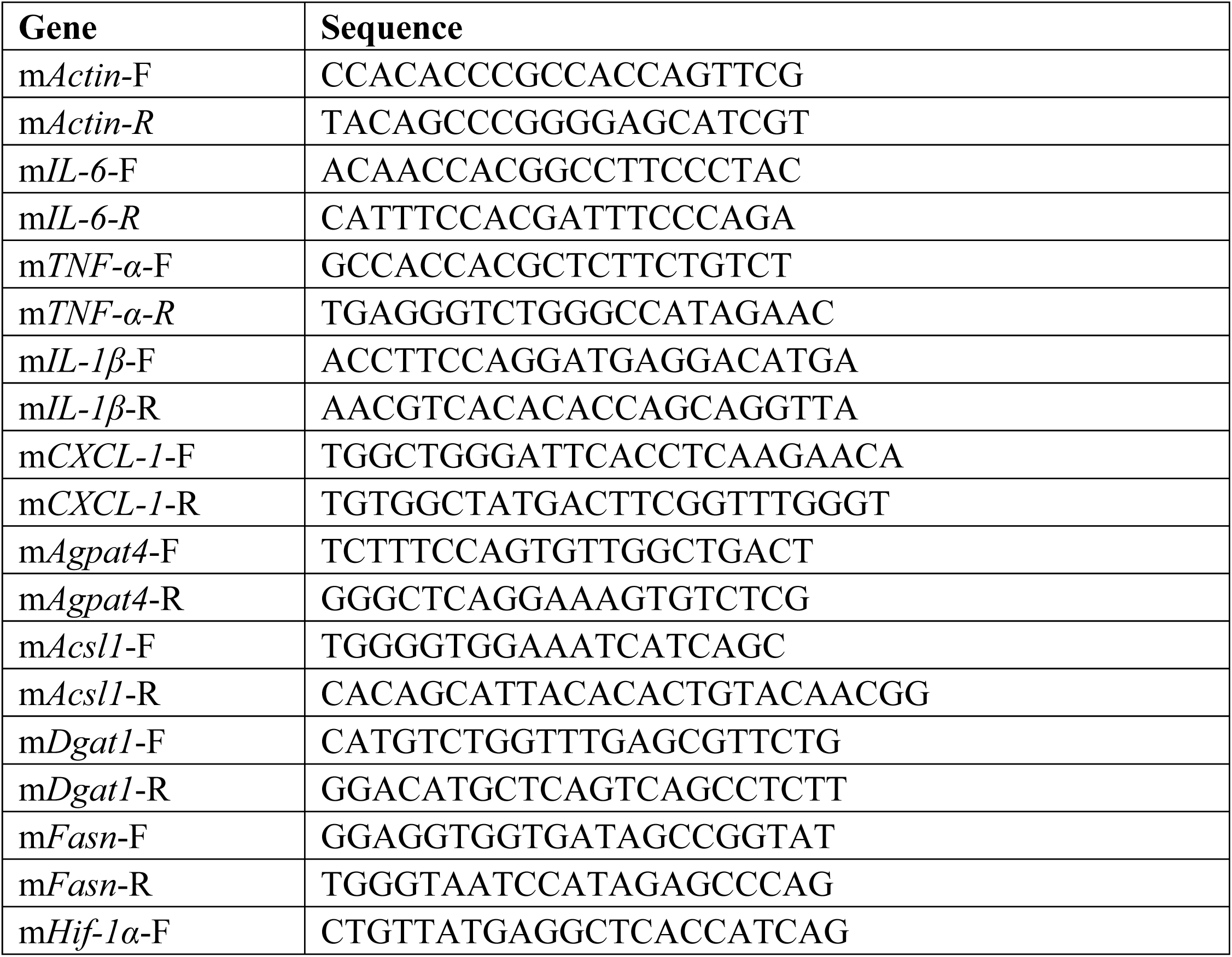
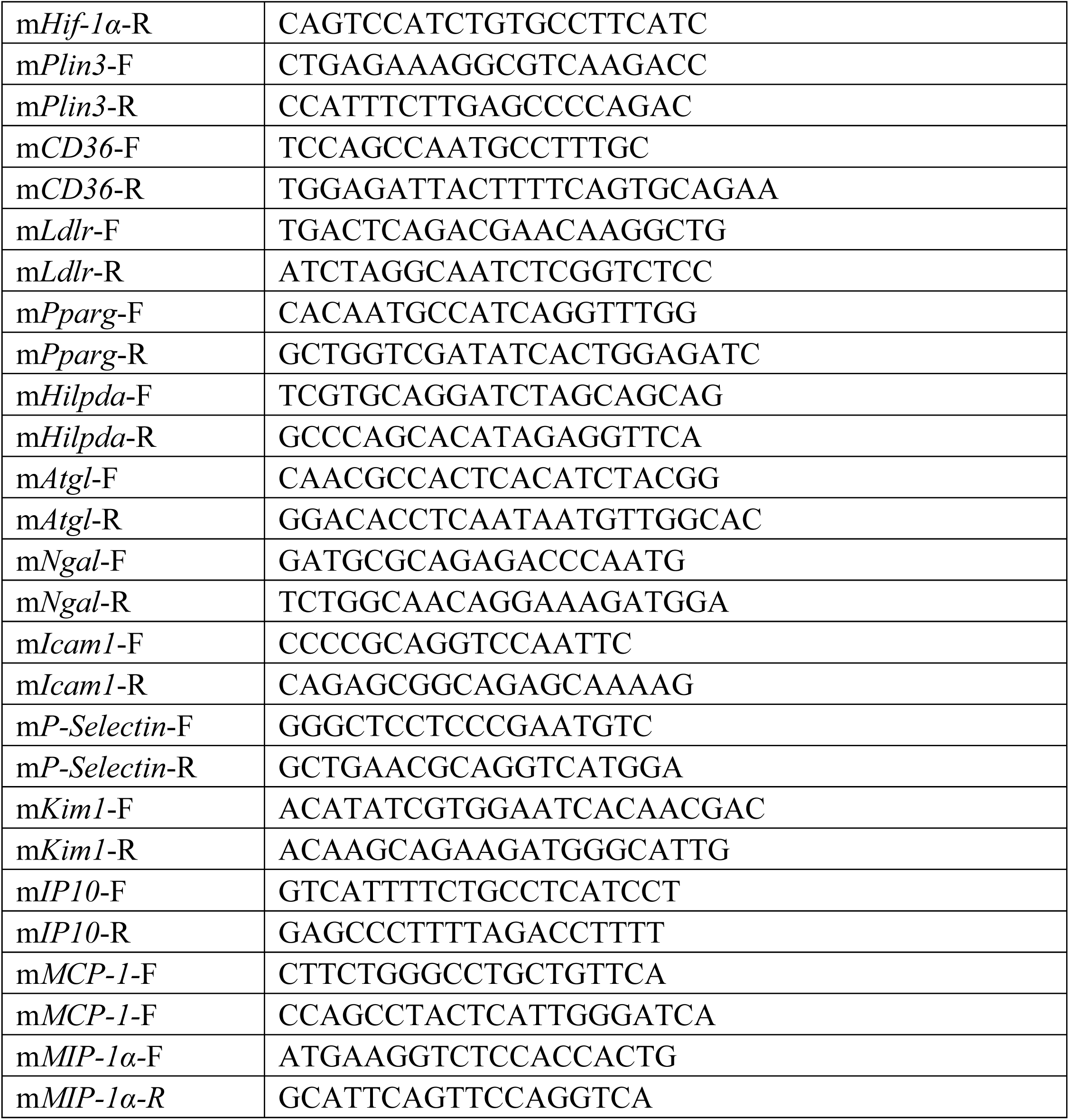
Primers for RT-qPCR.

