## Supplementary figures and images for "Lipid droplets restrict phagosome formation during *Candida* challenge"

### Supplemental Figure1

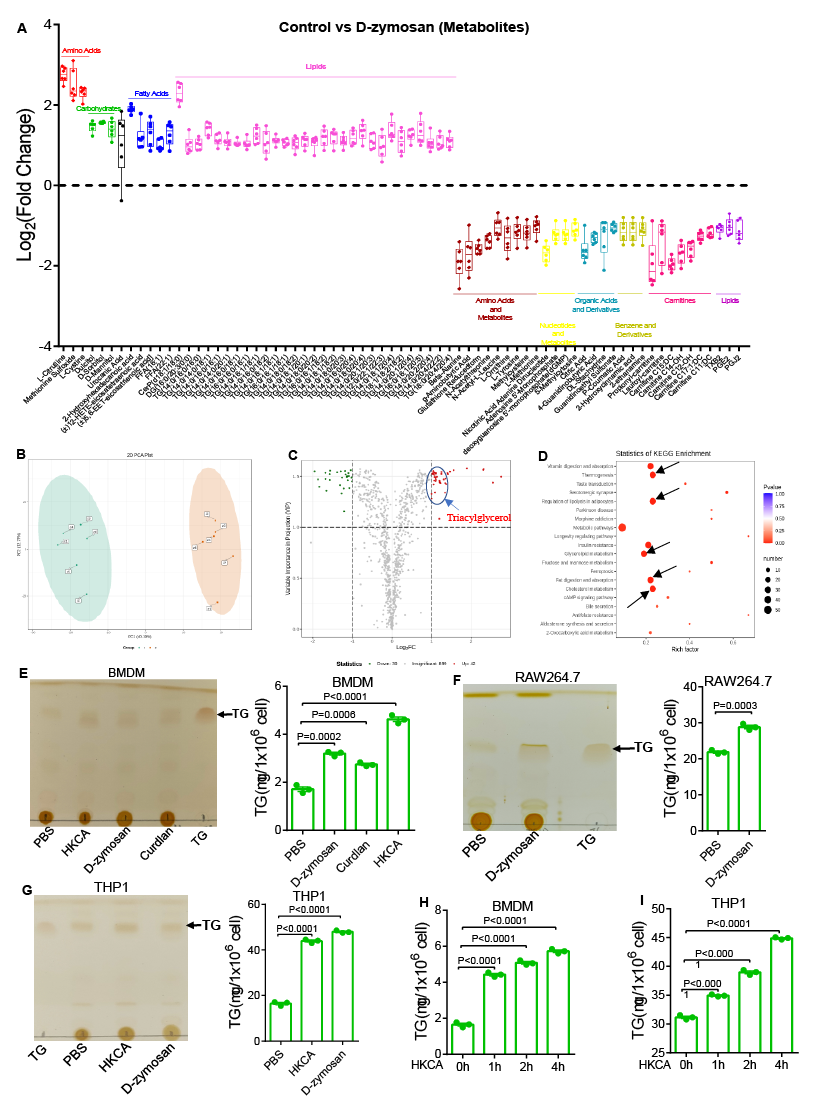

### Supplemental Figure2

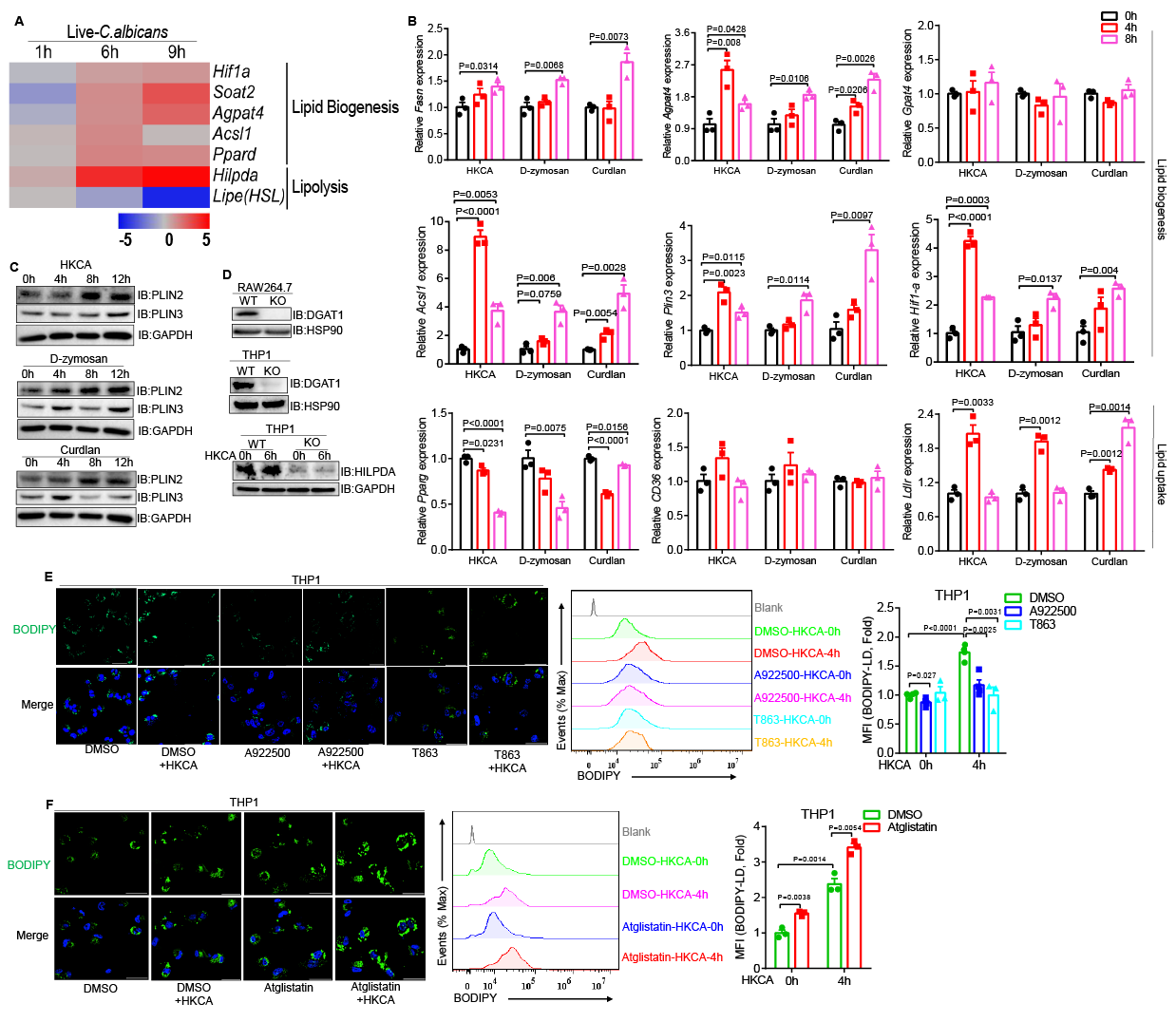

### Supplemental Figure3

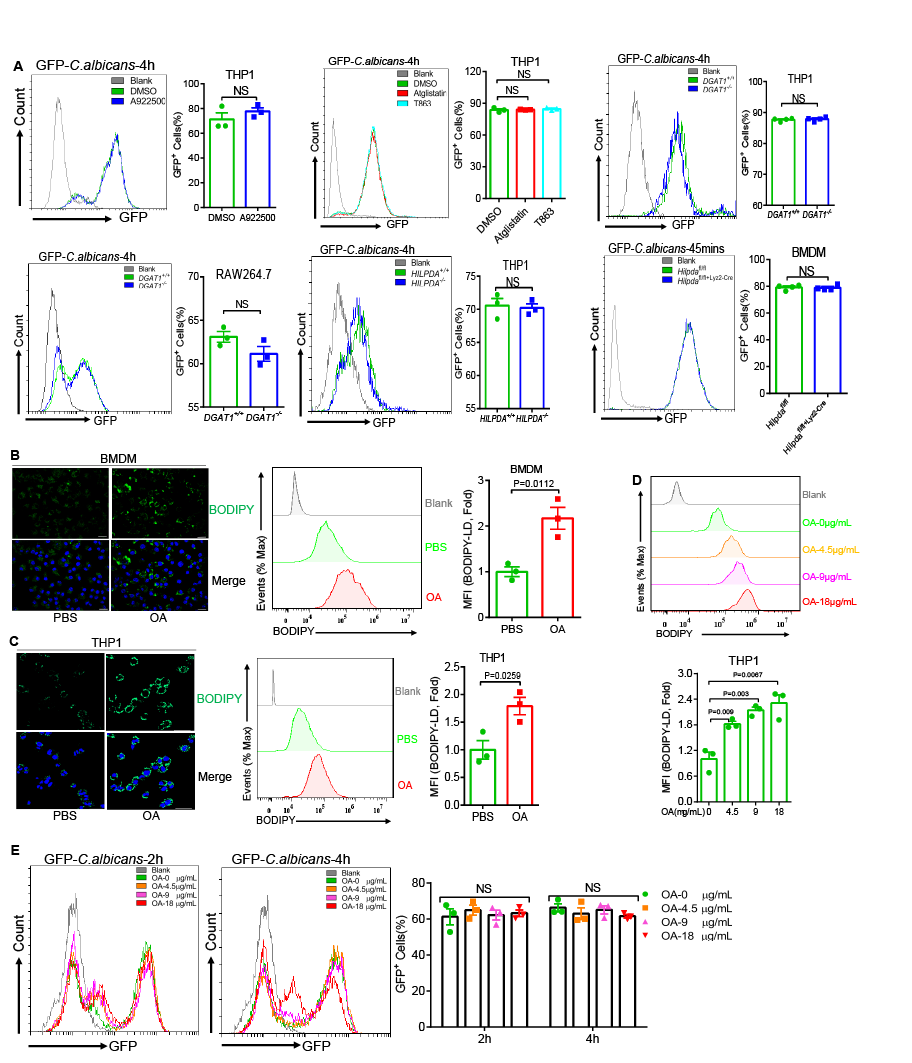

### Supplemental Figure4

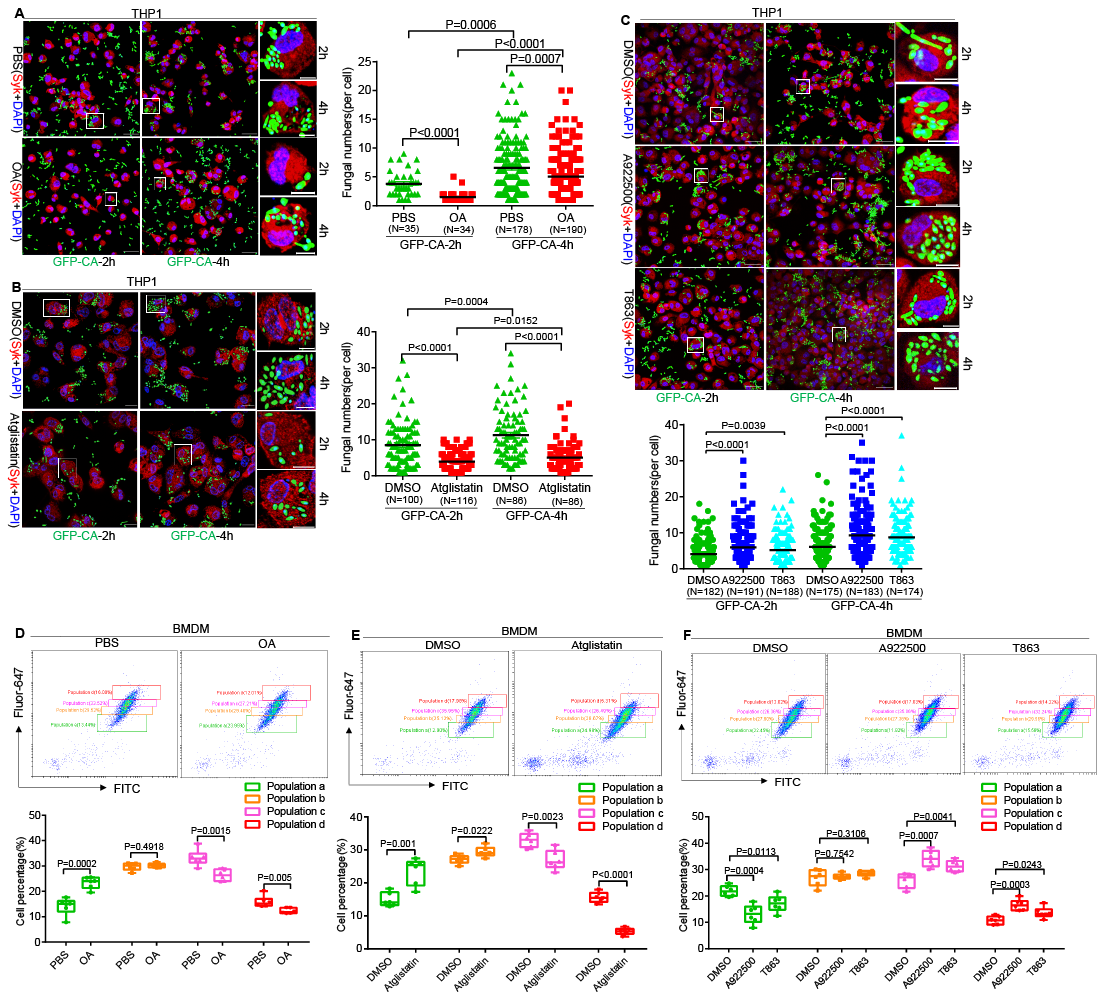

### Supplemental Figure5

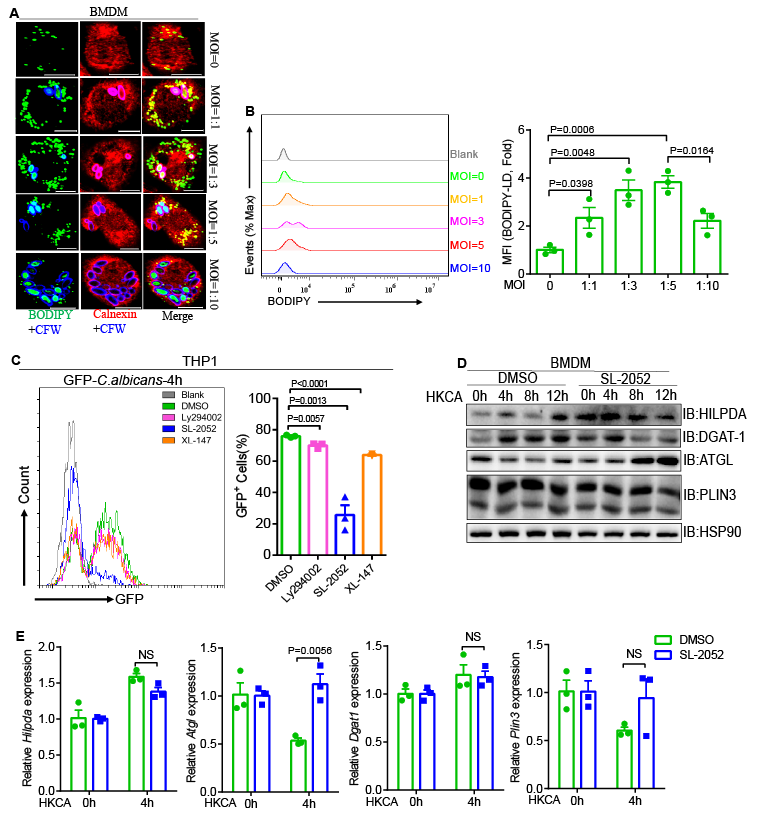

### Supplemental Figure6

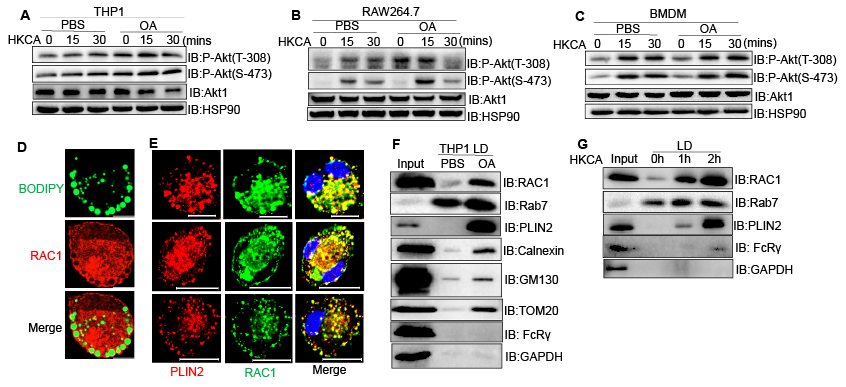

### Supplemental Figure7

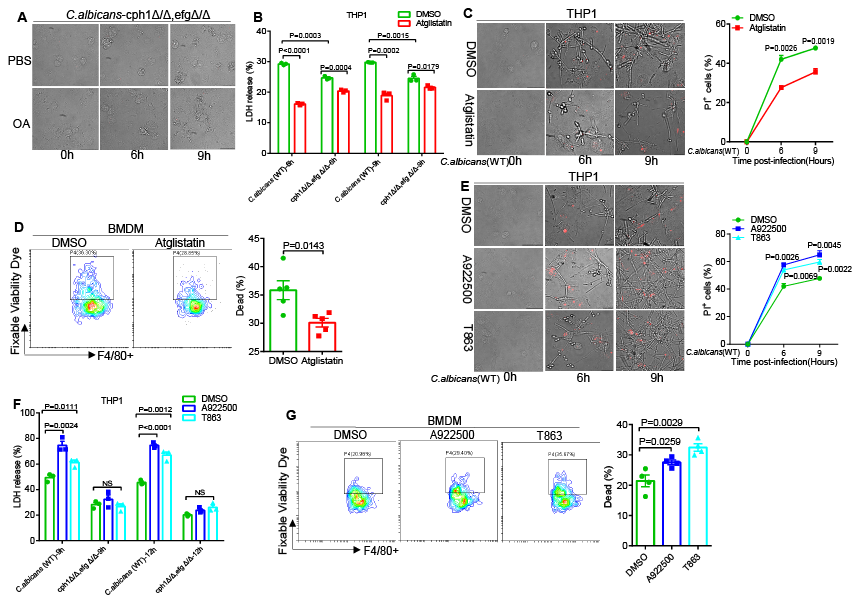

### Supplemental Figure8

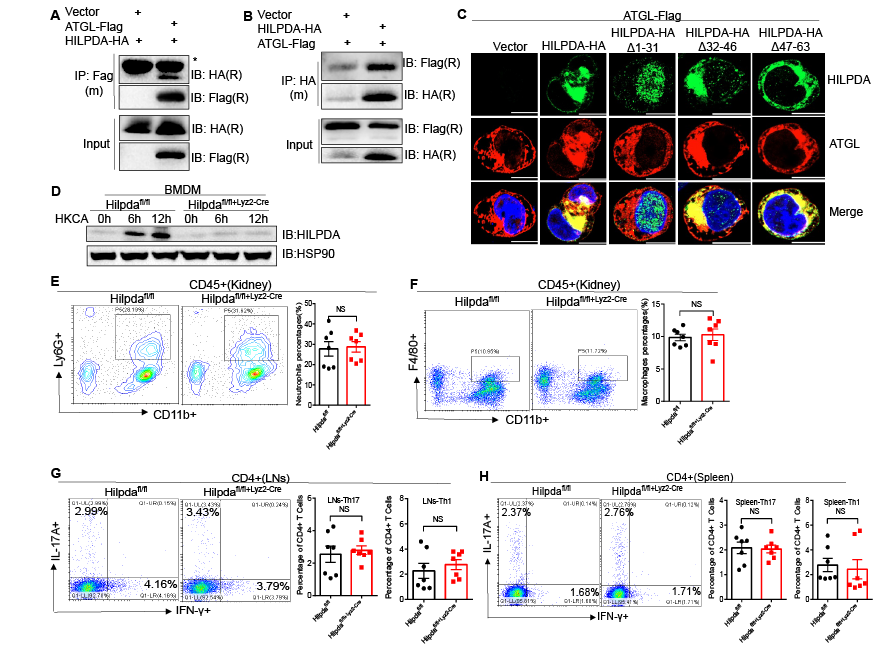

### Supplemental Figure9

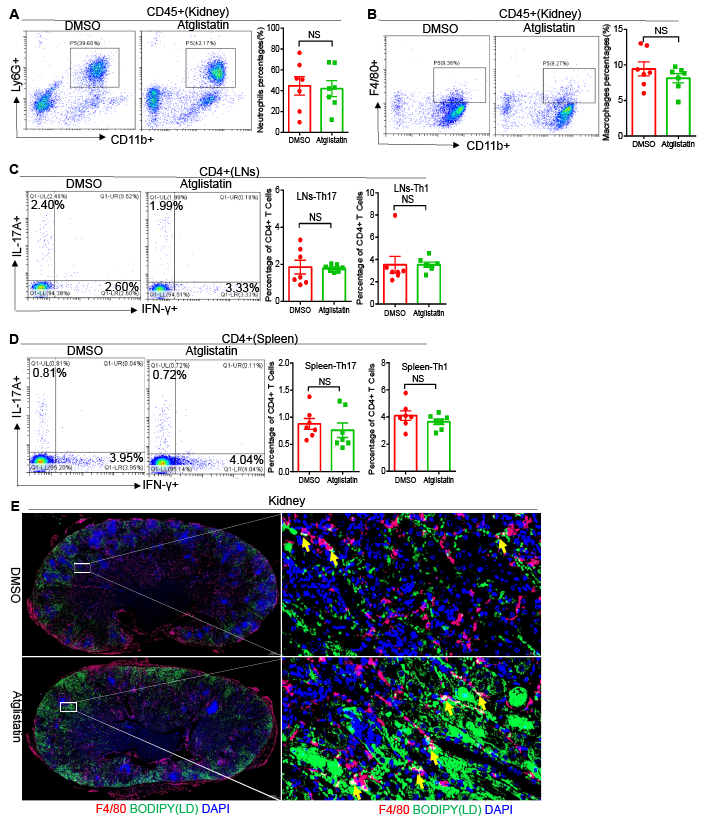

### Supplemental Figure10

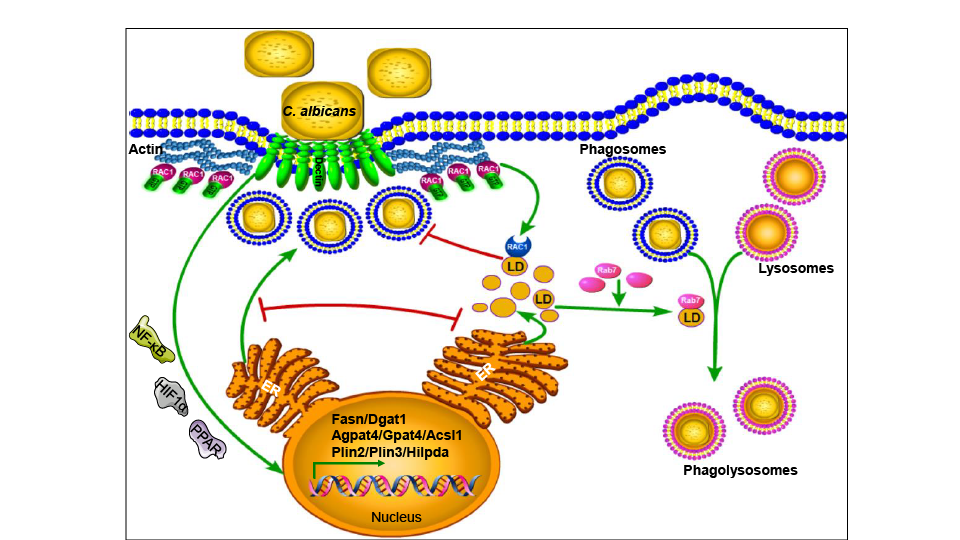
